# Chromatin activation as a unifying principle underlying pathogenic mechanisms in multiple myeloma

**DOI:** 10.1101/740027

**Authors:** Raquel Ordoñez, Marta Kulis, Nuria Russiñol, Vicente Chapaprieta, Renée Beekman, Cem Meydan, Martí Duran-Ferrer, Núria Verdaguer-Dot, Guillem Clot, Roser Vilarrasa-Blasi, Leire Garate, Estíbaliz Miranda, Arantxa Carrasco, Teresa Ezponda, Amaia Vilas-Zornoza, David Lara-Astiaso, Daphné Dupéré-Richer, Joost H.A. Martens, David Torrents, Halima El-Omri, Ruba Y Taha, Maria J. Calasanz, Bruno Paiva, Jesus San Miguel, Paul Flicek, Ivo Gut, Ari Melnick, Constantine S. Mitsiades, Jonathan D. Licht, Elias Campo, Hendrik G. Stunnenberg, Xabier Agirre, Felipe Prosper, Jose I. Martin-Subero

## Abstract

Multiple myeloma (MM) is a plasma cell neoplasm associated with a broad variety of genetic lesions. In spite of this genetic heterogeneity, MMs share a characteristic malignant phenotype whose underlying molecular basis remains poorly characterized. In the present study, we examined plasma cells from MM using a multi-epigenomics approach and demonstrated that when compared to normal B cells, malignant plasma cells showed an extensive activation of regulatory elements, in part affecting co-regulated adjacent genes. Among target genes upregulated by this process, we found members of the NOTCH, NFkB, mTOR1 signaling and p53 signaling pathways. Other activated genes included sets involved in osteoblast differentiation and response to oxidative stress, all of which have been shown to be associated with the MM phenotype and clinical behavior. We functionally characterized MM specific active distant enhancers controlling the expression of thioredoxin (*TXN*), a major regulator of cellular redox status, and in addition identified *PRDM5* as a novel essential gene for MM. Collectively our data indicates that aberrant chromatin activation is a unifying feature underlying the malignant plasma cell phenotype.

## INTRODUCTION

MM is an aggressive hematological neoplasm characterized by the uncontrolled expansion and accumulation of malignant plasma cells (PCs) in the bone marrow (San Miguel 2014; Kumar et al. 2017). Given the central role of genomic alterations in cancer development, the study of molecular mechanisms underlying MM pathogenesis has mostly focused on the genetic aberrations of these tumors (Stratton et al. 2009). Such studies have revealed that MM patients are genetically heterogeneous, without a single and unifying genetic event identified in all patients (Robiou du Pont et al. 2017), but nevertheless the central paradox of MM remains, i.e. terminally-differentiated plasma cells normally do not divide. This indicates that the essential aspects of the gene regulatory networks within the malignant plasma cell are dysfunctional and suggests that epigenetic deregulation of gene expression may be a root cause of the disease. Over the last decades, a greater understanding of the role of histone, DNA and other chromatin modifications in the epigenetic control of gene expression has transformed our understanding of transcriptional patterns in normal and neoplastic cells (Allis and Jenuwein 2016). In spite of the multifaceted nature of the epigenome (Stricker et al. 2017), the cancer epigenomics field has been mostly focused on the association between DNA methylation and transcription (Esteller 2008). However, recent whole-genome DNA methylation studies indicate that this association is less clear than previously appreciated (Kulis et al. 2012; Kretzmer et al. 2015; Kulis et al. 2015), and that understanding gene deregulation in cancer requires the integrative analysis of various epigenetic marks including histone modifications and chromatin accessibility (Martens et al. 2010; Ooi et al. 2016; Beekman et al. 2018). In MM, although several reports have identified alterations in the DNA methylome of malignant plasma cells (Walker et al. 2011; Heuck et al. 2013; Kaiser et al. 2013; Agirre et al. 2015), their pathogenic impact has been uncertain and the chromatin regulatory network underlying aberrant cellular functions in MM has just started to be characterized (Agarwal et al. 2016; Jin et al. 2018).

## RESULTS

### Integrative analysis of multiple epigenetic layers in MM

We performed an integrative analysis of multiple epigenetic layers in purified plasma cells from 3 MM patients, including whole-genome maps of six core histone modifications with non-overlapping functions (i.e. H3K27ac, H3K4me1, H3K4me3, H3K36me3, H3K27me3, and H3K9me3) by ChIP-seq, chromatin accessibility by ATAC-seq, DNA methylation by whole-genome bisulfite sequencing (WGBS) and gene expression by RNA-seq. The major limiting factor for such multi-omics characterization was the high amount of starting material needed from each individual patient. To overcome this limitation, we also analyzed the chromatin regulatory landscape of an extended series of MM patients, including additional profiles for H3K27ac (n=11), H3K4me1 (n=7), ATAC-seq (n=14) and RNA-seq data (n=37). Normal control consisted of reference epigenomes from B cell subpopulations at different maturation stages, including naive B cells (both from peripheral blood, pb-NBCs, and tonsils, t-NBCs), germinal center B cells (GCBCs), memory B cells (MBCs) and tonsillar PCs (t-PCs) (Beekman et al. 2018) (**Fig. 1a, Supplementary Tables 1 and 2**). Additionally, we were able to characterize the H3K27ac and transcriptional profiles of bone marrow plasma cells (bm-PCs), the normal counterpart of MM, from which a complete reference epigenome could not be generated due to the scarcity of this cell subpopulation in healthy bone marrow.

**Figure 1.**
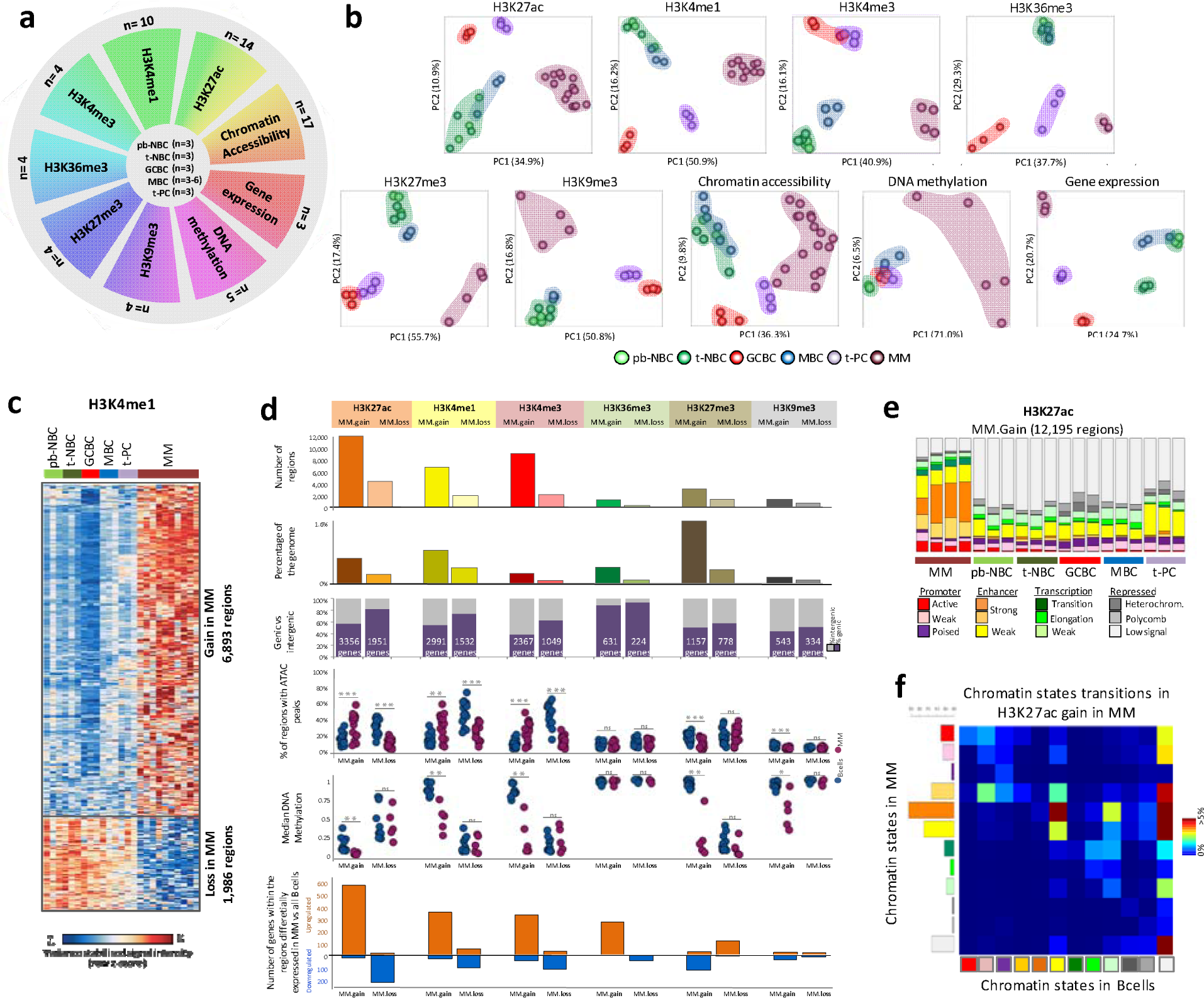
Initial characterization of epigenetic layers in multiple myeloma. a) Schematic representation of the experimental design. The outer circle shows the numbers of MM samples for the nine epigenetic layers used in the study, while numbers of normal B cell samples are shown in the centre. b) Unsupervised principal component analysis for the nine layers of the epigenome. c) Heatmap representation of the regions with differential H3K4me1 occupancy in MM as compared to a stable pattern throughout normal B cell differentiation. d) Characterization of regions with stable chromatin profiles throughout B cell differentiation showing either gain or loss of a specific histone mark in MM. From upper to lower panel: barplot showing number of regions detected for each condition; barplot showing total occupancy of the differential histone mark regions as a percentage of the whole genome; fractions of regions located in intergenic regions or inside genes (number of host genes associated to the differential regions shown in the graph); fraction of regions in MM (n=17) and normal B cells (n=15) harbouring ATAC-seq peaks within regions with increase or decrease of particular histone marks in MM; median DNA methylation levels in MM (n=5) and normal B cells (n=12) within the regions with increase or decrease of particular histone mark in MM; barplots presenting number of host genes associated with the differential histone mark regions that are upregulated or downregulated in MM as compared to normal B cells (FDR<0.05; |FC|>1.5). e) Distribution of the different chromatin states in all analyzed samples separately at regions with increase of H3K27ac in MM as compared to normal B cells. f) Chromatin state transition matrix for regions with increase of H3K27ac in MM as compared to normal B cells. Columns represent the chromatin state in normal B cells and rows are chromatin states in MMs that arise from normal B cells. The total matrix represents 100 percent of the differential regions. All p-values were calculated using a Student’s t-Test (two-sided). * p <0.05, ** p < 0.01, *** p < 0.001, ns: not significant. pb-NBC, naive B cells from blood; t-NBC, naive B cells from tonsils; GCBC, germinal center B cells; MBC, memory B cells; t-PC, plasma cell from tonsils; MM, multiple myeloma

Unsupervised principal component analysis indicated that MM displays a distinct epigenetic configuration than normal B cell subpopulations, which is reflected in every single layer of the epigenome (**Fig. 1b**). As B cell differentiation entails modulation of all studied epigenetic layers, we focused our analysis on the genome fraction changing in MM but showing stable chromatin profiles across the normal B cell maturation program. This strategy shall allow us to identify chromatin changes specifically altered during myelomagenesis (**Fig. 1c and Supplementary Fig. 1 and 2**). Overall, we observed that each histone mark undergoes more gains than losses in MM, but at different degrees, with H3K27ac, H3K4me1 and H3K4me3showing the largest number of gained regions (**Fig. 1d**). This finding points to a gain of regulatory elements such as enhancers and promoters in MM (**Fig. 1e and Supplementary Fig. 3**) that frequently arise from low-signal heterochromatic regions in normal B cells (**Fig.1f and Supplementary Fig. 4**). These regions were associated with more accessible chromatin, loss of DNA methylation and an increased expression of the associated genes in MM as compared to normal B cells (**Fig. 1d**). Loss of regulatory elements in MM was related to the opposite patterns with the exception of DNA methylation, which showed similar levels in normal B cells and MM in these regions. This finding indicates that once a region has become active and demethylated earlier in B cell differentiation, this methylation state is maintained regardless of changes in activity, thus retaining a memory of past activation, as previously described (Kulis et al. 2015; Beekman et al. 2018). MM-specific changes in H3K27me3 and H3K9me3 showed minor or absent modulation of chromatin accessibility and gene expression levels. In contrast, regions gaining these repressive marks in MM showed a marked loss of DNA methylation, which is counterintuitive, as these chromatin marks are conventionally known to be associated with methylated DNA (Vire et al. 2006; Schlesinger et al. 2007; Brinkman et al. 2012). We analyzed the DNA methylome of our MM samples in further detail and observed extensive DNA hypomethylation taking place in inactive chromatin regions regardless of the particular mode of repression, i.e. presence of H3K27me3, presence of H3K9me3 or low signal heterochromatin (**Fig. 1d and Supplementary Fig. 5a,b**). Remarkably, regions changing H3K27me3 or H3K9me3 frequently reflect transitions among repressive chromatin states from normal to neoplastic cells (**Supplementary Fig. 4**), and DNA methylation in MM seems to decrease in repressed chromatin independently of these transitions (**Supplementary Fig. 5c**). Collectively, these findings indicate that MM is characterized by a highly dynamic chromatin landscape, which on the one hand affects heterochromatin without apparent functional impact, and on the other hand, leads to an emergence of active regulatory elements leading to extensive perturbation of the MM transcriptome.

### *De novo* chromatin activation affects genes related to key pathogenic mechanisms in MM

To detect regulatory elements that could represent MM-specific epigenetic alterations, we aimed at identifying *de novo* active regions by focusing on H3K27ac, a *bona fide* indicator of enhancer and promoter activation (Zentner et al. 2011; Bonn et al. 2012). In this way, wecould select regions without any H3K27ac peak across normal B cell differentiation (i.e. inactive) while gaining this histone mark specifically in MM. Although due to the scarcity of plasma cells in the healthy bone marrow, it was not feasible to generate multiple epigenetic marks in this cellular subpopulation, we included bm-PC in this analysis as a key filter for H3K27ac and RNa-seq. We know that this information is limited but it has been very important in our study to discard changes associated to cell of origin (**Supplementary Fig. 6**). Using this strategy, we retained 1,556 regions, which were associated with 1,059 target genes, and were used for further downstream functional analyses (**Fig. 2a and Supplementary Table 3**). Within these 1,556 *de novo* active regions, we identified 806 sites that also increased chromatin accessibility in MM as compared to normal B cells. These sites were enriched in binding motifs of *IRF, FOX* and *MEF2* transcription factor (TF) families (**Fig. 2b and Supplementary Table 4**), which have been previously linked to MM pathogenesis (Carvajal-Vergara et al. 2005; Shaffer et al. 2008; Campbell et al. 2010; Bai et al. 2017; Agnarelli et al. 2018). In addition, we observed that some members of these families are overexpressed in MM as compared to normal PCs, such as *IRF1, IRF4, FOXO4, FOXP2, MEF2A* and *MEF2C* (**Fig. 2b and Supplementary Table 5**). Whereas part of them are *de novo* expressed in MM as compared to normal B cell differentiation, i.e. *IRF1* and *FOXP2*, others show elevated levels in normal PCs and are further upregulated in MM, i.e. IRF4 and *FOXO4*, and a third group is downregulated in normal PCs as compared to NBCs, GCBs and MBCs but showed increased expression in MM (**Fig. 2b**). These results support the hypothesis that extensive chromatin activation in MM may not only be mediated by disease-specific overexpression of TFs but also by exploiting and enhancing the function of TFs modulated during normal B cell differentiation.

Next, to decipher the downstream pathogenic relevance of *de novo* active regulatory regions in MM, we explored the functional categories associated with the target genes (**Fig. 2c**). The results point to a variety of functions previously described to be altered in MM (Bommert et al. 2006; Hu and Hu 2018), including regulation of osteoblast differentiation, multiple signaling pathways, such as NFkB signaling, mTOR1 signaling, p53 pathway and NOTCH pathway, as well as oxidative stress responses (**Fig. 2c**). Figure panels **2d-f** show examples of genes with regulatory elements becoming *de novo* active in MM as compared to normal cells. For instance, in the case of NOTCH pathway, we identified chromatin activation of genes at different steps of the pathway, starting from *de novo* active ligands, receptor processing machinery, up to downstream targets (**Fig. 2e**). As activation of the above-mentioned pathways is in part mediated by the crosstalk between MM and microenvironmental cells, our results suggest that such microenvironmental interactions may underlie chromatin activation of downstream genes in MM. Additionally, we observed a significant association with GO terms and hallmark signatures involved in oxidative stress responses (**Fig. 2f**). In fact, oncogenic transformation in MM is accompanied by higher endoplasmic reticulum and oxidative stresses mostly because of high immunoglobulin production and increased metabolic demands caused by the proliferative activity of MM cells (White-Gilbertson et al. 2013; Lipchick et al. 2016). Our findings suggest that MM cells activate a specific regulatory network to be able to survive in spite of elevated oxidative stress conditions. This network seems to be related to the emergence of *de novo* enhancers leading to overexpression of genes involved in major detoxification systems, including e.g. *GLRX, OXR1, ATOX1, PYROXD2* and thioredoxin (*TXN*) (**Fig. 2f and Fig. 3a**).

**Figure 2.**
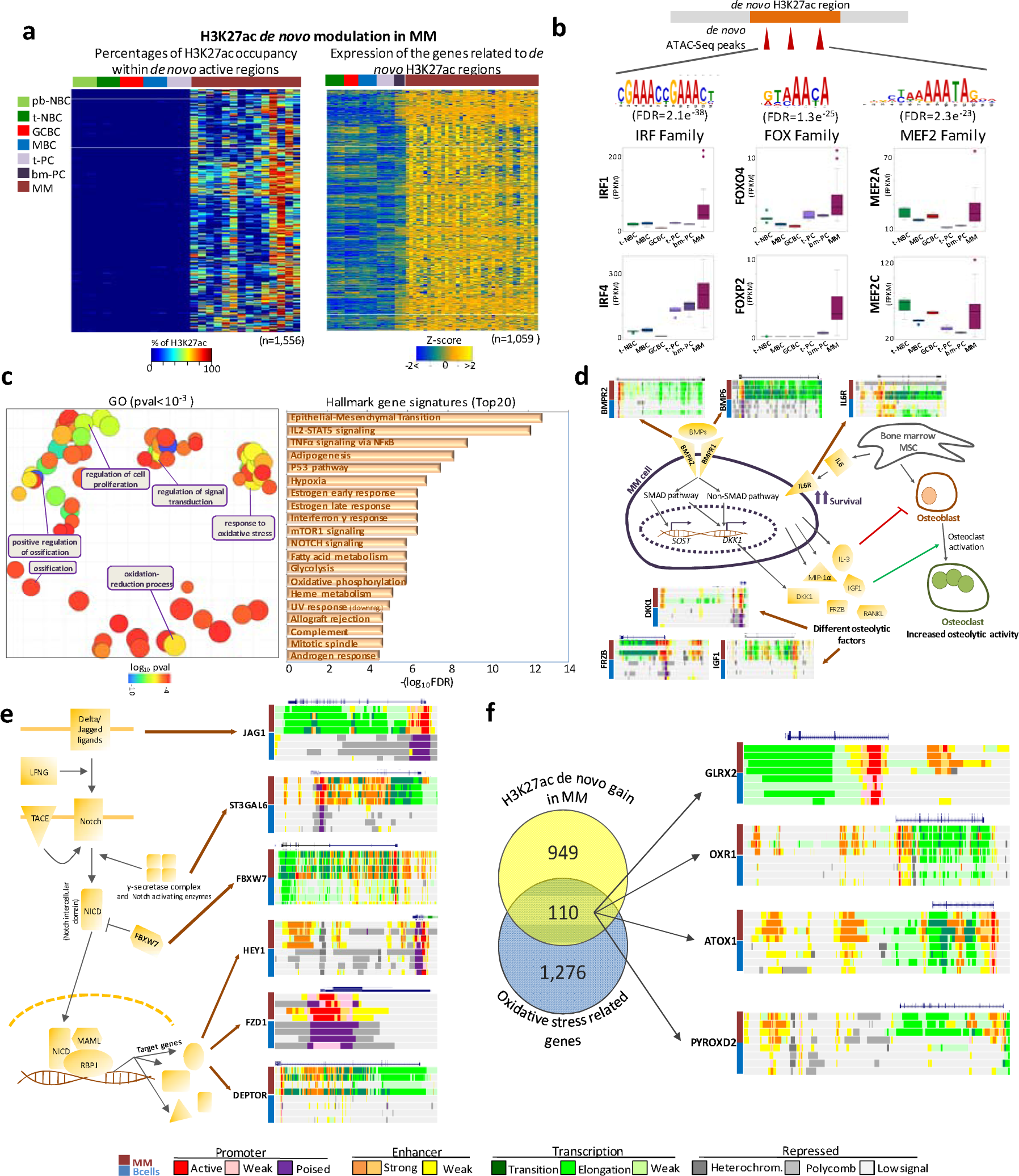
Functional impact of *de novo* chromatin activation in MM. a) Heatmaps representing the percentage of the regions covered byH3K27ac peak within the *de novo* activated regions in MM (left panel) and expression of the genes associated with these regions. b) TF families motifs enriched in the *de novo* activated regions in MM, i.e. IRF, FOX and MEF2, as identified by MEME analysis. For each TF family, expression of two selected members, upregulated in MM as compared to normal B cells is shown. c) Gene ontology results, shown as semantic-similarity scatter plot of the most significant GO terms (p<10-3), summarized together by ReVigo software (left panel) and a list of top 20 hallmark gene signatures, determined using MSigBD Collection (right panel). d) Schematic representation of mechanisms involved in interactions between MM and the bone marrow microenvironment, with selected genes harboring activated chromatin in MM as compared to normal B cells. e) Schematic representation of the NOTCH signaling pathway with selected genes harboring activated chromatin in MM as compared to normal B cells. f) Venn Diagram presenting the overlap of genes associated with *de novo* active regions in MM and genes belonging to GOs related with oxidative stress (i.e. oxidative-reduction process and response to oxidative stress). Chromatin states within selected genes in MM and normal B cells are shown on the right panel. pb-NBC, naive B cells from blood; t-NBC, naive B cells from tonsils; GCBC, germinal center B cells; MBC, memory B cells; t-PC, plasma cell from tonsils; bm-PC, plasma cell from bone marrow; MM, multiple myeloma.

### *TXN de novo* activated enhancer is an essential regulatory element in MM

*TXN* contributes to maintain reactive oxygen species (ROS) homeostasis in the cell to prevent oxidative damage (Arner and Holmgren 2006) and has been recently shown to enhance cell growth in MM and its inhibition leads to ROS-induced apoptosis in MM cell lines (Raninga et al. 2015; Sebastian et al. 2017). From the chromatin perspective, the *TXN* gene body and promoter show activating histone modifications both in MM and normal B cell subpopulations. Remarkably, we identified a *de novo* active distant enhancer region, located approximately 50 kb upstream *TXN* transcription start site (TSS), which may account for elevated activity of this gene (**Fig. 3a**). This overall active chromatin was associated with *TXN* overexpression in both MM patients and cell lines compared to normal B cell differentiation (**Fig. 3b and Supplementary Fig. 7a**). We further aimed to characterize the pathogenic implication of *TXN* deregulation in MM cells. *TXN* inactivation using the CRISPR/Cas9 strategy in MM cell lines (**Fig. 3c,d and Supplementary Fig. 7b,c**) led to a significant reduction of growth rate as compared to control cells, as well as an increased apoptotic phenotype, as measured by Annexin V flow cytometry analysis (**Fig. 3e,f and Supplementary Fig. 7d,e**). These results corroborate that *TXN* is an essential gene for MM cell survival and proliferatio (Raninga et al. 2015; Raninga et al. 2016; Zheng et al. 2018). We then studied whether *TXN* overexpression was driven by the *de novo* activation of the distant enhancer identified in MM patients. We designed a CRISPR/Cas9 paired gRNAs system, with two plasmids carrying gRNAs flanking the regulatory region, to completely delete the 11kb *TXN* enhancer identified in MM patients (**Fig. 3a**). We observed that enhancer deletion in bulk cells significantly reduced *TXN* expression at the RNA and protein level in a MM cell line and was associated with a significant decrease in cell proliferation (**Fig. 3g-j and Supplementary Fig. 8**). Taken together, our results extend previous reports on the essential role of *TXN* in MM and show that *de novo* activation of a distant enhancer underlies its overexpression and pathogenic impact.

**Figure 3.**
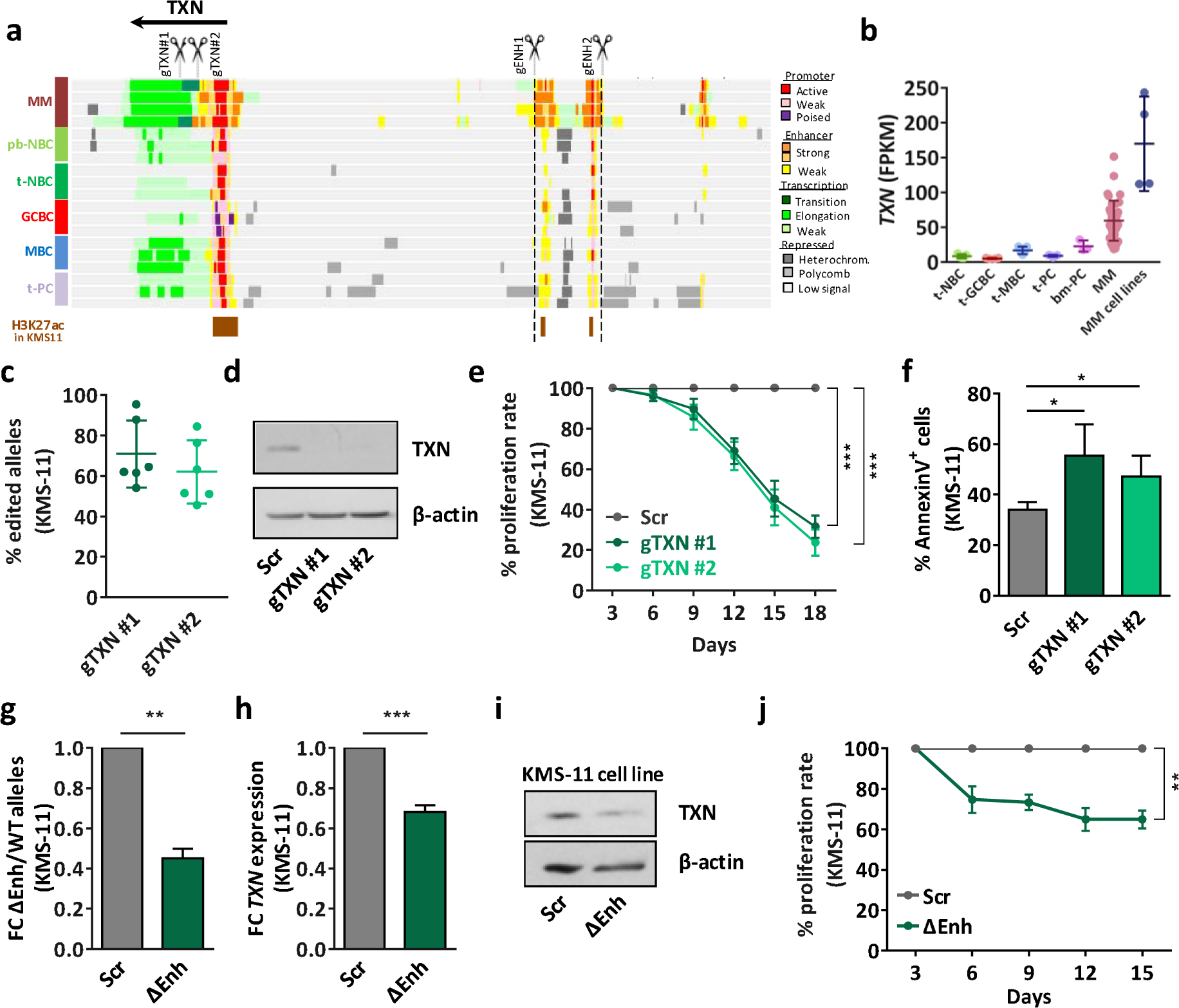
TXN de novo activated enhancer as an essential regulatory element in MM. a) Genome browser snapshot of the TXN locus and the associated enhancer de novo activated in MM located at 50 Kb from the promoter region. Displayed tracks represent the chromatin state annotation in MM patients and normal B cells and an additional track of H3K27ac peaks in KMS-11 cell line (ENCODE Consortium, ENCSR715JBO). gRNA design strategy for TXN knockout and enhancer deletion is also shown. b) TXN expression in MM patients and cell lines analyzed by RNA-seq. c) Estimation of the allelic cell population percentage exhibiting indel events in the targeted site, analyzed by Tracking of Indels by DEcomposition (TIDe) web tool. d) Validation of TXN knockout by western blot analysis in KMS-11 cell line. e) Cell proliferation assay comparing the growth proliferation rates of scramble cells (Scr) and cells harboring two different gRNAs, as determined by flow cytometry analysis. f) Effect of TXN knockout in cell apoptosis, as determined by Annexin V flow cytometry analysis. g) Quantification of TXN enhancer deletion by genomic DNA qPCR normalized to a distal non-targeted genomic region, represented as fold change of deleted enhancer (ΔEnh) versus wild type (WT) alleles. h) TXN mRNA expression levels determined by RT-qPCR. i) TXN protein levels in cells with deleted enhancer region (ΔEnh) and scrambled cells (Scr) determined by western blot. j) Cell proliferation assay comparing the growth proliferation rates of scramble cells and cells harboring the enhancer deletion (ΔEnh), as determined by flow cytometry analysis. * p<0.05, ** p<0.01, *** p<0.001, ns: not significant. pb-NBC, naive B cells from blood; t-NBC, naive B cells from tonsils; GCBC, germinal center B cells; MBC, memory B cells; t-PC, plasma cell from tonsils; bm-PC, plasma cell from bone marrow; MM, multiple myeloma.

### *PRDM5* as a new candidate oncogene in MM

Beyond the activation of genes affecting relevant pathogenic mechanisms in MM, we observed that adjacent genes without evident functional association become *de novo* active in MM. This suggests the presence of co-regulated chromatin regions that can coordinate the deregulation of more than one target genes specifically in tumor plasma cells (**Fig. 4a**). To capture these co-regulated chromatin regions, we selected adjacent co-expressed genes (pearson R>0.5, p<0.05) from the list of 1,059 genes with *de novo* chromatin activation using RNA-seq data from 37MM samples. In this way, we detected 42 pairs and one triplet of co-expressed genes (**Supplementary Table 6**). From this list, we selected one co-regulated chromatin region containing two unrelated genes, *PRDM5*, a TF member of the PRDM family that in contrast to *PRDM1* is not expressed in normal plasma cells but is reported to act as a tumor suppressor gene in solid tumors (Deng and Huang 2004; Duan et al. 2007; Watanabe et al. 2007; Shu et al. 2011); and *NDNF*, a gene involved in neuronbiology (Kuang et al. 2010; Ohashi et al. 2014). These two co-expressed genes reside in a 500 kb region showing clear *de novo* activation in MM (**Fig. 4a,b**), and such co-expression was validated in an additional sample cohort of 10 MM patients and 10 MM cell lines by RT-qPCR (**Supplementary Fig. 9a**). In addition to their co-expression, both genes were clearly upregulated in MM, with negligible expression levels across B cell differentiation, including t-PCs and bm-PCs (**Fig. 4b**). Next, we studied whether *PRDM5* and *NDNF* co-activation is related with a specific three-dimensional genome organization of the region. We performed 4C-seq experiment in a MM cell line expressing both transcripts (U266) and a mantle cell lymphoma cell line (JVM-2) as negative control. We identified three-dimensional interactions between *PRDM5* and *NDNF* gene loci only in U266 MM cells, suggesting that the topological remodeling of this region in MM may be related to the coordinated activation of both genes (**Fig. 4c**). Finally, in order to identify whether these physically-related, but functionally-independent genes are involved in MM pathogenesis, we performed doxycycline-inducible shRNA mediated knockdown in different MM cell lines. Gene knockdown at the transcriptional and protein level was validated by RT-qPCR (**Fig. 4d and Supplementary Fig. 9b**) and western blot (**Supplementary Fig. 9e**) to ensure the correct silencing of the target gene. *PRDM5* knockdown induced a significant decrease in cell proliferation rate and increased cell death (**Fig. 4e,f and Supplementary Fig. 9c,d**). However, we did not detect any significant effect in cell viability after *NDNF* silencing (**Fig. 4e,f and Supplementary Fig. 9c,d**). Taken together, these results suggest that this *de novo* activation of the co-regulated chromatin region comprising *PRDM5* and *NDNF* drives the coordinated overexpression of both genes but only *PRDM5* seems to be related to MM pathogenesis. This evidence infers that *PRDM5* acts as an oncogene in MM, which contrasts with its previously reported tumor suppressor role in solid tumors (Duan et al. 2007; Shu et al. 2011; Shu 2015). In order to decipher the specific function of *PRDM5* in MM pathogenesis, we performed RNA-seq analysis comparing the transcriptional landscape of *PRDM5* silenced vs. mock MM cell line. We found 1,216 differentially expressed genes (**Fig. 4g**). Those genes downregulated upon *PRDM5* silencing were associated with the cell cycle as well as various types of cellular stress responses such as DNA damage stress induced by UV light and unfolded protein response stress, suggesting that *PRDM5* may be involved in protecting MM cells against the cellular stress created by proliferation and high protein synthesis (**Fig. 4h**).

**Figure 4.**
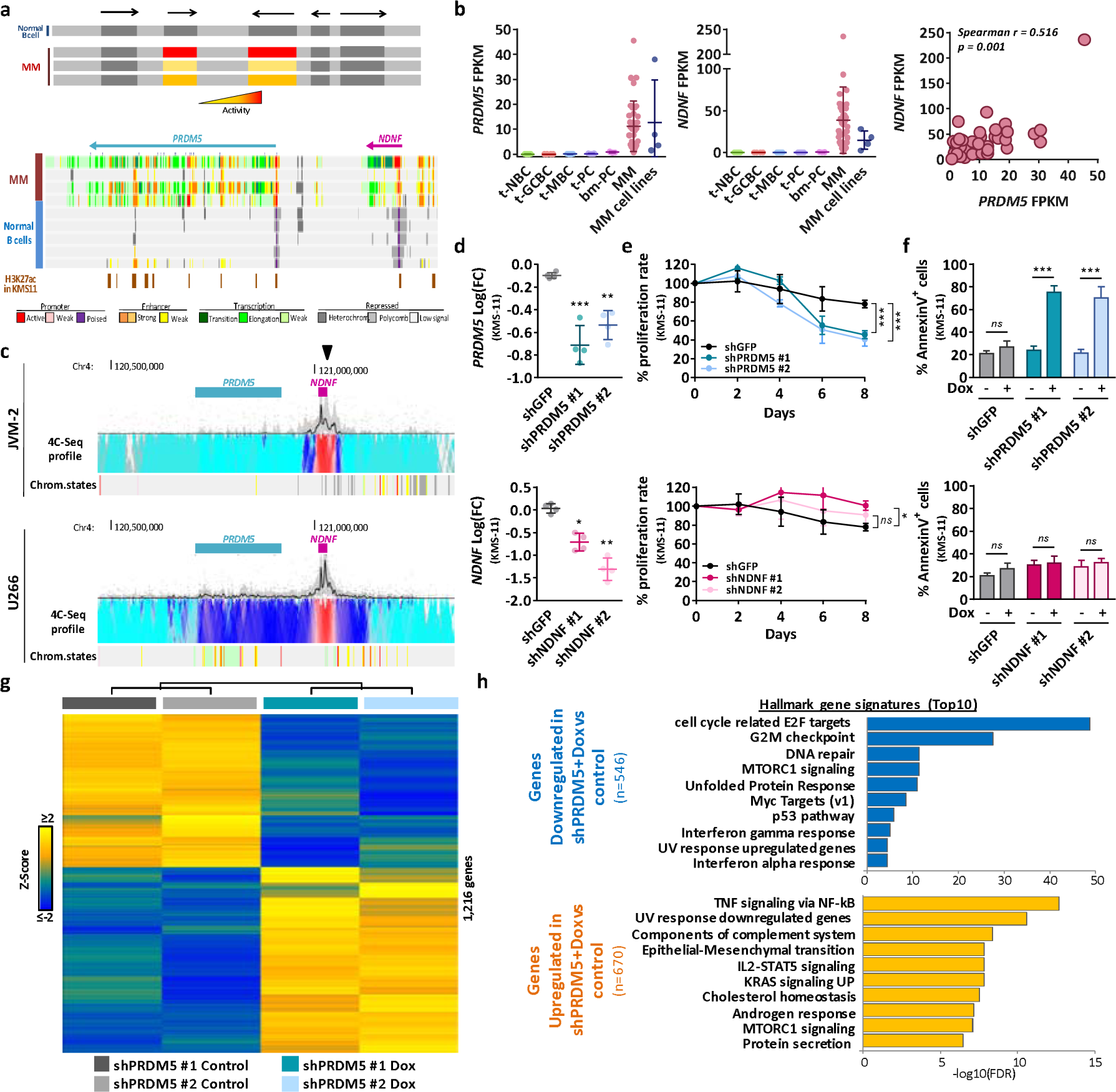
Identification of PRDM5 as a new candidate oncogene in MM. a) Schematic strategy for identification of co-regulated chromatin regions. Genome browser snapshot of the co-regulated chromatin region comprising the PRDM5 and NDNF locus. Displayed tracks represent the chromatin state annotation in MM patients and normal B cells, and an additional track of H3K27ac peaks in KMS-11 cell line (ENCODE Consortium, ENCSR715JBO). b) PRDM5 and NDNF expression analyzed by RNA-seq (left and center panel), and correlated expression levels in MM patients (right panel). c) Normalized levels of chromatin interaction frequencies from the indicated viewpoint (black arrowhead) between PRDM5 and NDNF gene loci as analyzed by 4C-seq in the JVM-2 mantle lymphoma cell line lacking PRDM5 and NDNF expression, and the U266 MM cell line expressing both transcripts. Lowest contact frequencies are indicated in turquoise, and highest frequencies in red. PRDM5 and NDNF location are shown above 4C-seq track, while chromatin states for each of these cell lines are shown below. d) Validation of PRDM5 (upper panel) and NDNF (lower panel) knockdown after shRNA expression determined by RT-qPCR. Expression values are normalized to the same condition prior to addition of doxycycline. Statistical analysis compares the effect of each shRNA versus the scramble shRNA (shGFP). e) Relative cell proliferation rate (%) of KMS-11 cell line normalized to the same condition prior to addition of doxycycline. The upper panel presents the results for PRDM5knockdown cells and the lower panel for NDNF knockdown cells. Statistical analysis compares the effect of each shRNA versus the scramble shRNA (shGFP). f) Effect of PRDM5 (upper panel) and NDNF (lower panel) knockdown in cell apoptosis, as determined by Annexin V flow cytometry analysis. g) Heatmap of gene expression levels (indicated as Z-scores) of differentially expressed genes in PRDM5 knockdown cells before and after 4 days of the addition of doxycycline (n = 2 for control and Dox groups; 1,216 genes). h) List of top 10 hallmark gene signatures, determined using MSigBD Collection for genes downregulated (upper panel) or upregulated (lower panel) in PRDM5 knockdown cells 4 days after the addition of doxycycline. Dox: Doxycycline. * p<0.05, ** p<0.01, *** p<0.001, ns: not significant. pb-NBC, naive B cells from blood; t-NBC, naive B cells from tonsils; GCBC, germinal center B cells; MBC, memory B cells; t-PC, plasma cell from tonsils; bm-PC, plasma cell from bone marrow; MM, multiple myeloma.

## DISCUSSION

Overall, we provide a multi-omics view of the MM epigenome in the context of normal B cell differentiation which extends recent efforts to analyze polycomb-mediated gene repression and regulatory networks driven by super-enhancer elements in MM pathogenesis (Agarwal et al. 2016; Jin et al. 2018). Our series contains cases from different genetic groups, reflecting the heterogeneous nature of the disease, however the sample size is not informative to characterize such biological variability. Considering such limitation, we decided to focus on identifying common epigenetic events involved in MM pathogenesis by integrating two data sets: first, a multilayered epigenomic characterization of MM patients that provided a global view of the chromatin function and deregulation; second, an extended series focused on the profiling of the chromatin regulatory landscape and the identification of regulatory elements. By the comprehensive integration of both data series, we could identify common MM-specific signatures for multiple epigenetic marks, revealing the existence of a core epigenomic landscape underlying MM pathogenesis, remarkably associated with the widespread activation of regulatory elements silenced not only in PCs, the normal counterpart of MM, but across the entire B cell maturation program. Such activation seems to be mediated by the action of specific TF families, such as IRF, FOX or MEF2, which are part of a TF network deregulated in MM (Jin et al. 2018). In addition, members of these three TF families have been previously reported to be functionally important for MM pathogenesis (Carvajal-Vergara et al. 2005; Shaffer et al. 2008; Campbell et al. 2010; Bai et al. 2017; Agnarelli et al. 2018). Thus, inhibition of these TFs may represent a rational therapeutic approach to revert aberrant chromatin activation in MM. This hypothesis is supported by previous publications indicating that IRF4 or FOXM1 inhibition induces cell death and influences the proliferation in MM cell lines, although efficient strategies to block these TF in MM patients are still missing (Shaffer et al. 2008; Morelli et al. 2015; Gu et al. 2016).

The altered chromatin landscape in MM affects genes involved in a variety of signaling pathways and cellular responses previously reported to play a central role into myelomagenesis (Bommert et al. 2006; Hu and Hu 2018). A remarkable feature of these findings is that MM cells *de novo* activate regulatory elements of genes involved in preventing cell death associated with oxidative stress. One of the major systems maintaining cellular redox homeostasis is the thioredoxin system (Arner and Holmgren 2006). We show that downregulation of *TXN* expression either by disrupting the gene itself or by deleting its distant regulatory element impair MM cell growth, representing a potential therapeutic target. In addition to the emergence of enhancer elements, we also identified the activation of co-regulated chromatin regions, which coordinate the deregulation of more than one adjacent target gene. We functionally analyzed the impact of two of these co-expressed genes, *PRDM5* and *NDNF*, and only the former has an impact on MM cell growth. Furthermore, through the transcriptional analysis of PRDM5 knocked-down MM cells, we discovered that this gene seems to be involved incellular responses to DNA damage and protein synthesis stresses. Indeed, a previous report has identified that *PRDM5* is upregulated upon UV-light exposure and its promoter contains binding sites of stress-related TFs (Shu et al. 2011). This finding further supports the concept that protection against various cellular stresses is a key element for the survival of MM cells, and that neoplastic plasma cells may in part achieve this stress-resistant phenotype through aberrant *de novo* chromatin activation. In conclusion, extensive chromatin activation seems to be a unifying principle underlying multiple pathogenic mechanisms in MM, which suggests that epigenetic drugs such as e.g. BET inhibitors may be appropriate as a backbone treatment for patients affected with this aggressive disease.

## ACKNOWLEDGEMENTS

This research was funded by the European Union’s Seventh Framework Programme through the Blueprint Consortium (grant agreement 282510), Fundació La Marató de TV3, Instituto de Salud Carlos III (ISCIII) PI14/01867, PI16/02024 and PI17/00701, TRASCAN (EPICA), MINECO Explora (RTHALMY), Departamento de Salud del Gobierno de Navarra 40/2016, Gilead Fellowship Program (GLD16/00142), Multiple Myeloma Research Foundation Networks of excellence, the International Myeloma Foundation (Brian van Novis) and the Qatar National Research Fund award 7-916-3-237. Furthermore, the authors would like to thank the support of the Generalitat de Catalunya Suport Grups de Recerca AGAUR 2017-SGR-736 and 2017-SGR-1142, TRASCAN-iMMunocelland European Research Council starting grant (MYELOMANEXT), CIBERONC (CB16/12/00489, CB16/12/00369 and CB16/12/00225), co-financed with FEDER funds, the Accelerator award CRUK/AIRC/AECC joint funder-partnership as well as NCI R01 CA180475 and a MMRF collaborative grant. R.O was supported by a FPU Fellowship of the Spanish Government, M.K. by an AOI grant of the Spanish Association Against Cancer and N.R. by the Acció instrumental d’incorporació de scientífics i tecnòlegs PERIS 2016 from the Generalitat de Catalunya. This work was partially developed at the Centro Esther Koplowitz (CEK, Barcelona, Spain). We particularly acknowledge the patients for their participation and the Biobank of the University of Navarra for its collaboration.

## AUTHOR CONTRIBUTIONS

Investigator contributions were as follows: H.E.-O., R.Y.T., M.J.C., B.P., J.S.M., X.A., and F.P. contributed to sample collection as well as to their biological and clinical annotation; M.K., N.R., N.V-D., R.V.-B., J.H.A.M., I.G., A.M., E.C., H.G.S., and J.I.M.-S. performed, coordinated and/or supported histone mark, ATAC-seq, 4C-seq, methylome and transcriptome data generation; R.O., M.K., V.C., R.B., C.M., M.D.-F., G.C., D.T., P.F., X.A., and J.I.M-S performed, coordinated and/or supported histone mark, ATAC-seq, 4C-seq, methylome and transcriptome data analysis; R.O., L.G., E.M., A.C., T.E., D.D.-R., C.S.M., J.D.L., X.A. and F.P. performed and/or supervised in vitro functional assays; R.O. M.K., C.S.M., J.D.L., E.C., H.G.S., X.A., F.P. and J.I.M.-S. participated in the study design and/or data interpretation. X.A., F.P. and J.I.M.-S. directed the research and wrote the manuscript together with R.O. and M.K.

