## Supplementary material for "Chromatin activation as a unifying principle underlying pathogenic mechanisms in multiple myeloma": Methods

**Collection and preparation of patient and normal samples.** Tumor samples used for epigenomic analyses were obtained from bone marrow aspirations of newly diagnosed patients of MM before administration of any treatment. All MM samples were purified by $CD138^+$ positive magnetic separation using AutoMACS system (AutoMACS Pro Separator, Miltenyi Biotech), obtaining over 90% purity in all cases, as assessed by flow cytometry. The data from normal B cells (i.e. pb-NBC, t-NBC, GCBC, MBC, t-PC) was generated by our lab previously, using samples collected and isolated as previously described[8,11]. Purified plasma cells (bm-PC) were selected from bone marrow aspirations from healthy donors ranging from 20 to 30 years. After Ficoll-Isopaque density gradient centrifugation, we performed a selective depletion of $CD3^+$, $CD14^+$ and $CD15^+$ cells by immunomagnetic selection (Miltenyi Biotec), followed by FACS of $CD45^+$ / $CD138^+$ / $CD38^+$ cells using a FACSAriaII (BD Biosciences) device (**Supplementary Table 1**).

For downstream transcriptional analyses, an additional series of MM cases and normal B cell controls was studied. Tumor samples were obtained and purified following the same strategy as described above, while naive B cells (NBC), germinal center B cells (GCBC), memory (MBC) and tonsillar plasma cells (t-PC) were isolated from human tonsils of healthy donors while bone marrow plasma cells (bm-PC) from bone marrow of healthy donors by multiparameter fluorescence-activated cell sorting (FACS) using the expression level of nine different surface antigens as previously described[48]. The following monoclonal antibody (MoAb) combination was used for the cell isolation from tonsils: CD45-OC515 (Clone HI30, Immunostep, Salamanca, Spain); CD20-Pacific Blue (Clone 2H7, Bioloegend, San Diego, California, UnitedStates); CD44-APCH7 (Clone G44-26, Beckton Dickinson, Dur-ham, North Carolina, United States). CD10 PE-Cy7 (Clone HI10a BecktonDickinson, Durham, North Carolina, United States); CD38-FITC (Clone LD38, Cytognos, Salamanca, Spain); CXCR4-PE (Clone 12G5, Beckton Dickinson, Durham, North Carolina, UnitedStates); CD27-APC (Clone L128, Beckton Dickinson, Durham, North Carolina, UnitedStates) and CD3-PerCP-Cy5.5 (Clone SK7, Beckton Dickinson, Durham, North Carolina, United States). bm-PC were sorted using FACS AriaIIfrom human bone marrow of healthy donors using CD38-FITC (Clone LD38, Cytognos, Salamanca, Spain); CD138-BV421 (Clone MI15, Beckton Dickinson, Dur-ham, North Carolina, United States) and CD27-BV510 (Clone 0323, Bioloegend, San Diego, California, UnitedStates) (**Supplementary Table 1**).

All patients and donors gave informed consent for their participation in this study, which was approved by the clinical research ethics committee of Clínica Universidad de Navarra.

**ChIP-seq, ATAC-seq, RNA-seq and WGBS data generation**. ChIP-seq of the six different histone marks and ATAC-seq data were generated as described (http://www.blueprint-epigenome.eu/index.cfm?p=7BF8A4B6-F4FE-861A-2AD57A08D63D0B58). Catalog numbers of antibodies (Diagenode) used are H3K27ac: C15410196/pAb-196-050 (LOT: A1723-0041D), H3K4me1: C15410194/pAb-194-050 (LOT: A1863-001P), H3K4me3: C15410003-50/pAb-003-050 (LOT: A5051-001P), H3K36me3: C15410192/(pAb-192-050 (LOT: A1847-001P), H3K9me3: C15410193/pAb-193-050 (LOT: A1671-001P), H3K27me3: C15410195/pAb-195-050 (LOT: A1811-001P).

Strand specific RNA-seq data of the reference epigenomes and the validation series was generated as previously described[49]. Briefly, RNA was extracted using TRIZOL (Life Technologies) and libraries were prepared using a TruSeq Stranded Total RNA Kit with Ribo-Zero Gold (Illumina). Adapter-ligated libraries were amplified and sequenced using 100bp single-end reads for the reference epigenome samples and 50bp paired-end reads for the samples in the validation series.

RNA-seq data for the shPRDM5 cell lines series was performed following MARS-seq protocol adapted for bulk RNA-seq[50] with minor modifications. Briefly, 150,000 cells were harvested in 100 ul of Lysis/Binding Buffer (Ambion), vortexed and stored at -80ºC until further processing. Poly-A RNA was reverse-transcribed using poly-dT oligos carrying a 7 bp-index. Pooled samples were subjected to linear amplification by IVT. Resulting RNA was fragmented and dephosphorylated. Ligation of partial Illumina adaptor sequences was followed by a second RT reaction. Full Illumina adaptor sequences were added during final library amplification. RNA-seq libraries quantification was done with Qubit 3.0 Fluorometer (Life Technologies) and size profiles examination with Agilent's 4200 TapeStationSystem. Libraries were sequenced in an Illumina NextSeq 500 at a sequence depth of 10 million reads per sample.

WGBS of the reference epigenomes was generated as previously described[8]. Briefly, 1–2 µg of DNA was sheared and fragments of 150–300 bp were selected using AMPure XP beads (Agencourt Bioscience). After adaptor ligation (Illumina TruSeq Sample Preparation kit), DNA was treated with sodium bisulfite using the EpiTexy Bisulfite kit (Qiagen). Two rounds of bisulfite conversion were performed to ensure a conversion rate of over 99%. Enrichment for adaptor-ligated DNA was carried out through seven PCR cycles and paired-end DNA sequencing (2 × 100 bp) was then performed using the Illumina HiSeq 2000 platform.

**Read mapping and initial data processing**. Fastq files of ChIP-seq data were aligned to genome build GRCh38 (using bwa 0.7.7, picard 2.8.1 and samtools 1.3.1) and wiggle plots were generated (using PhantomPeakQualTools 1.1.0) as described (http://dcc.blueprint-epigenome.eu/#/md/methods).

Peaks of the histone mark data were called as described (http://dcc.blueprint-epigenome.eu/#/md/methods) using MACS2 (version 2.1.1.20160309), with input control. ATAC-seq fastq files were aligned to genome build GRCh38 using bwa 0.7.759 (parameters: -q 5, -P, -a 480) and SAMTOOLS v1.3.160 (default settings). BAM files were sorted and duplicates were marked using PICARD tools v2.8.1 (http://broadinstitute.github.io/picard, default settings). Finally, low quality and duplicates reads were removed using SAMTOOLS v1.3.160 (parameters: -b, -F 4, -q 5, -b, -F 1024). ATAC-seq peaks were determined using MACS2 (v2.1.1.20160309, parameters: -g hs -q 0.05 --keep-dup all -f BAM –nomodel –shift -96 –extsize 200) without input control. For downstream analysis peaks with p-values <1e-3 (H3K36me3, H3K9me3 and H3K27me3) or <1e-7 (H3K4me3, H3K4me1, H3K27ac, ATAC-seq) were included. For each mark a set of consensus peaks, only including regions on chromosome 1-22, present in the normal B cells (n=15 for histone marks and n=18 for ATAC-seq) and in the MM samples (n=4 for H3K4me3, H3K36me3, H3K9me3, H3K27me3; n=14 for the H3K27ac series, n=10 for the H3K4me1 series and n=17 for the ATAC-seq series) was generated by merging the locations of the separate peaks per individual sample. To generate the consensus peak file for the reference epigenomes, only peaks with input were used except for ATAC-seq for which peaks without input were used. For ChIP-seq, the numbers of reads per sample per consensus peak were calculated using the genomecov function of bedtools. For the ATAC-seq, the number of insertions of the Tn5 transposase per sample per consensus peak were calculated by first determining the estimated insertion sites (shifting the start of the first mate 4bp downstream), followed by the genomecov function of bedtools. Using DESeq2[53], variance stabilizing transformation (vst) values were calculated for all consensus peaks. The number of consensus peaks per chromatin mark were 72,848 (H3K4me3), 60,494 (H3K4me1), 125,026 (H3K27ac), 33,907 (H3K36me3), 61,142 (H3K9me3), 34,499 (H3K27me3), and 121,860 (ATAC-seq). Principal component analyses (PCA) were generated with the prcomp function in R using the (corrected) vst values of all peaks that were present in >1 sample, thus eliminating possible individual-specific peaks.

ChIP-seq of H3K27ac in bm-PC was processed as described above and peaks were called as described (http://dcc.blueprint-epigenome.eu/#/md/methods) using MACS2 (version 2.1.1.20160309).

Three different variants of RNA-seq were performed, which were processed as follows: 1) RNA-seq data of the reference epigenome samples (3 MM and 15 normal B cells) was aligned to genome build GRCh38, signal files were produced and gene quantifications (gencode 22;19,736 protein coding genes) were calculated as described (http://dcc.blueprint-epigenome.eu/#/md/methods) using the GRAPE2 pipeline with the STAR-RSEM profile (adapted from the ENCODE Long RNA-seq pipeline). The expected counts and FPKM estimates were used for downstream analysis. The PCA of the RNA-seq data was generated with the prcomp function in R using log10 transformed FPKM (+0.01 pseudocount) data. 2) RNA-seq of additional validation series (37 MM and 22 normal B cells) was aligned in a two-step procedure, using STAR v2.4 to remove any potential ribosomal RNA leftover from the ribodepletion step. A first pass alignment was done against human ribosomal sequences allowing multi-mapping. Unmapped reads from the first step were then aligned to hg19 human reference using GENCODE v19 junction points. Cufflinks v2.2.1 was run on the resulting alignment files to create de novo transcriptome assembly specific to each sample using strand-specific settings and GENCODE v19 as a database. Cufflinks outputs for all of the samples were then merged with themselves and GENCODE database using cuffmerge. R and GNU parallel was used to facilitate the analysis. 3) Low input 3' end RNA-seq data of shPRDM5 knockdown cells was aligned to the GRCh38/hg38 human genome with STAR version 2.5.2B with default parameters. To assign reads to genes we applied feature Counts.

Mapping and determination of methylation estimates was performed using GEMBS, as described by authors[51]. We retained 9,214,561 CpGs (chr1-22) showing at least 10 reads in all 20 samples. All downstream analyses were performed with this DNA methylation matrix.

**Detection of differential epigenetic regions and *de novo* active regions.** For individual histone marks and ATAC-seq data, only consensus regions present in at least 1 and in a maximum of all but 1 sample were used, thus excluding individual specific and constitutive regions. Of the included consensus peaks, those differentials among MM as compared to normal B cell subpopulations were defined using the likelihood ratio test (FDR < 0.01) of the DEseq2 package. Next, we separated the regions with either stable or dynamic patterns throughout normal B cell differentiation. For that purpose, we performed differential analysis within normal B cell subpopulations by DEseq2 package, classifying those with FDR<0.01 as regions harbouring epigenetic mark modulation throughout normal B cell maturation, and the regions with FDR>0.01 as well as those that have no peaks at all in normal B cells as regions with stable chromatin during B cell maturation. The latter list was used to identify regions with either gain or loss of a particular epigenetic mark in MM (**Supplementary Fig. 1**).

To stringently identify regions *de novo* active in MM, several filters were applied to the previously detected set of regions with differential gain of H3K27ac (n=12,195). This strategy is depicted in **Supplementary Figure 6**. First, the H3K27ac profile of bm-PC from one healthy donor was generated and the peaks shared between MM and bm-PC were filtered out. Next, the percentages of H3K27ac peak occupancy within each differential region were calculated and those regions having less than 20% of H3K27ac in normal B cells while showing a consistently present H3K27ac gain in MM (FDR<0.05) were selected. Finally, in order to identify the regions with an impact on gene expression, we performed the following steps: 1) *de novo* H3K27ac regions were annotated to all the genes lying within the same topological associated domain (TAD, data from lymphoblastoid cell line, GM12878); 2) gene expression changes between MM and normal B cells were identified by comparing RNA-seq data from 37 MM cases and 22 normal mature B cell subtypes samples, including 3 also healthy bm-PCs (data from additional validation RNA-seq series); and 3) only the genes overexpressed in MM as compared to all normal B cells (Limma, FDR<0.05) and with clearly higher expression in MM versus bm-PC (|FC| >1.5) were selected. This strategy led to the identification of 1,059 target genes corresponding to 1,556 regions with *de novo* gain of H3K27ac in MM. The identify regions and target genes de novo repressed in MM, the same type of analysis was performed with all the analogous steps but for the contrary situation. In this case, only 4 repressed regions were identified, corresponding to 6 genes downregulated in MM.

**Detection of differentially methylated CpGs.** Detection of differentially methylated CpGs (DMCs) in MM as compared to normal B cells was performed using limma with methylation difference between conditions of 0.25 and FDR <0.05. Furthermore, detection of DNA methylation valleys (DMVs) was performed as described by Wei Xieetal[52]. Briefly, we employed a window-based approach to determine DMVs. To identify each DMVs in each sample, the genome was first divided in 1kb bins and the DNA methylation level was averaged within each bin. Then a sliding 5kb window (with 1kb step) was used to identify regions that have an median methylation level less than 0.15 in a 5kb window in MM samples. Continuous regions resulting from this analysis were then merged to form DMVs in each sample.

**Differential expression analysis.** In this study, three different types of RNA-seq were analyzed, depending on the purpose of the study, i.e. RNA-seq of the reference epigenome samples, RNA-seq of additional validation series and low input RNA-seq for characterization of transcriptional changes in samples from PRDM5 knockdown studies. For the first two RNA-seq experiments, Limma analysis was performed in order to obtain differentially expressed protein coding genes (FDR < 0.05, |FC|>1.5) between MM samples and all B cell subpopulations. The threshold to consider a gene as unexpressed was set to 0.1 FPKM. For the low input 3' end RNA-seq dataset of shPRDM5 knockdown cells, only genes that were expressed (sum of FPKM values in all samples greater than 10) were included. Differential expressed genes were defined using the likelihood ratio test of DEseq2 package (|FC|> 1.5 and p < 0.01).

**Gene ontology and canonical pathways analysis of data sets.** GO enrichment was performed using GOrilla online tool[54], and further summarized together by ReVigo software[55] in order to represent them as semantic-similarity scatter plot of the most significant GO terms. Additionally, hallmark gen set analysis was performed, using MSigDB Collections platform[56,57].

**Transcription factor analysis**. Enrichment for TF binding sites was analyzed in regions gaining chromatin accessibility within the 1,556 *de novo* active regions in MM. Gain of ATAC-seq peaks was determined as regions without any ATAC-peak in the normal B cells (n=15) while showing a significant gain of chromatin accessibility in the MM cohort (n=17; FDR<0.01). Using this strategy, we ended up with 806 sites. Enrichment analysis of known TF binding motifs was performed using the AME tool from MEME suite[58], from the non-redundant homo sapiens 2018 Jaspar database (537 TF motifs), applying a one-tailed Wilcoxon rank-sum test with the maximum score of the sequence, a 0.05 FDR cutoff and a background formed by reference GRCh38 sequences extracted from the consensus ATAC-seq peaks enriched in at least two samples.

**Cell culture**. KMS-11, RPMI8226, MM.1S and U266 MM cell lines, as well as JVM-2 cell line were maintained in RPMI-1640 medium (Lonza) supplemented with 20% FBS (Gibco), 1%

Penicillin / Streptomycin (Lonza) and 2% HEPES (Life Technologies). All cells were maintained at 37 $^oC$ and 5% $CO_2$. HEK293T cells were maintained in DMEM (Lonza) with high glucose and pyruvate, supplemented with 10% FBS (Gibco), 1% Penicillin / Streptomicyn (Lonza) and 2% HEPES (Life Technologies) and grown in a humidified atmosphere at 37 $^oC$ and 5% $CO_2$. Only cells at low passage (below passage 16) were used for lentiviral production.

**4C-seq data generation and analysis.** 4C templates for JVM-2 and U266 cell lines were prepared as previously described[59,60], using $10^7$ cells per 4C library. We performed this experiment for the *NDNF* promoter (chr4: 121,070,660-121,071,025) using DpnII and BfaI as first and second restriction enzymes and the following respective primers: 5'-TTGCTTCTCATCTGTCGATC-3' and 5'-CAGAAAGGTGAACCGAGAG-3'. Data analysis was performed with the 4C-seq pipe pipeline using default settings and removing reads corresponding to self-ligated or non-digested fragments.

**Inducible shRNA knockdown system and lentiviral production.** Short hairpin RNAs (shRNA) against each of the genes of interest (i.e. *PRDM5* or *NDNF*) were designed using public available algorithms (http://www.broad.mit.edu/science/projects/rnai-consortium/trc-shrna-design-process and https://www.med.nagoya-u.ac.jp/neurogenetics/i_Score/i_score.html). shRNAs were cloned into a TET inducible version of pLKO.1 vector (Tet-pLKO-puro, Addgene #21915) after EcoRI/AgeI digestion. A shRNA targeting GFP gene (shGFP) was used as experimental control. See **Supplemental Table 7** for target sequences and sequencing primers.

Lentiviruses were generated by co-transfecting HEK293T cells with shRNA-encoding plasmid, psPAX2 (Addgene, #12260) and pMD2G (Addgene, #12259) plasmids in a 3:2:1 ratio, using Lipofectamine 2000 (Invitrogen). Growth media was exchanged after 12 hours. Lentivirus-containing supernatant was harvested 48 hours later, filtered (0.2 µm) and applied directly to cells for infection with 2 µg/ml polybrene (Sigma). Target cell lines were selected in 1.5 µg/ml puromycin (Sigma) for 72 hours.

Stable cell lines harboring the silenced shRNA-expressing cassette were grown in Tet-Free FBS (Fetal Bovine Serum (FBS) South America, Tetracycline Free, Biowest) conditions to prevent Tet-promoter basal expression. shRNA expression was induced by adding 1 µg/ml doxycycline (Sigma) to the culturing media and replaced every 48 hours. Target gene knockdown was validated by RT-qPCR to ensure the correct silencing of the target gene.

***TXN* editing by CRISPR-Cas9 system.** To perform CRISPR-Cas9 editing of *TXN* gene, we first generated two stable Cas9 cell lines: KMS-11 and MM.1S, infecting them with lentiviruses carrying the Cas9-2A-Blasticidin expressing cassette (Lenti-Cas9-2A-Blast plasmid, Addgene #73310) following standard protocol, described before, and selecting them with 10 µg/ml blasticidin (Invitrogen) for 7 days. For *TXN* knockout, we designed two different gRNAs (gTXN#1 and gTXN#2), both targeting exon 2 of the gene, and we cloned them into two separetedly in CRISPseq-BFP-backbone (gift from Ido Amit laboratory, Addgene #85707). See **Supplementary Table 7** for target sequences and sequencing primers. In case of genomic deletion of *TXN* 11kb enhancer region, paired gRNAs flanking the region of interest were designed and cloned separately: one into CRISPseq-BFP-backbone harboring a BFP reporter gene and the other into pLKO5.sgRNA.EFS.GFP (Addgene #57822) harboring a GFP reporter. See **Supplementary Table 7** for target sequences. Corresponding lentiviruses were produced for all the constructs (as described above), and constitutively Cas9 expressing cell lines were infected. $BFP^+$ cells in the case of single gRNAs and double $BFP^+$ / $GFP^+$ cells in the case of double gRNAs were determined by flow cytometry (FACS Canto II, BD Biosciences) 72 hours after infection. Percentage of $BFP^+$ cells and $BFP^+$ / $GFP^+$ exceeded 90% in all cases (**Supplementary Fig. 8a**). As a control, cells infected with empty CRISPseq-BFP-backbone and/or pLKO5.sgRNA.EFS.GFP were used (scramble, Scr).

To assess editing efficiency in the TXN knockout experiment, genomic DNA was extracted 72h after infection from the total pull of cells, targeted loci were amplified and sequenced by Sanger reaction (for primer sequences see **Supplementary Table 7**). The spectrum and frequency of targeted mutations generated in the cell pool was determined by the public available software TIDE[61]. In the case of TXN enhancer deletion, excision was assessed qualitatively by PCR using primers flanking the genomic target region, such that the edited alleles should produce a shorter PCR product than the uncut alleles in the total pull of cells, 72h after infection (**Supplementary Fig. 8b**). For a more quantitative determination of the editing efficiency in the total pull of cells, qPCR was performed using a primer internal to the deleted region, which will only amplify the wild type allele (**Supplementary Table 7**). For statistical analysis, normality test for each group were performed using Shapiro-Wilk. Student t-test was used for parametric group comparisons, whereas Mann-Whitney test was used for non-parametric comparisons. Protein levels of target genes were assessed by western blot analyses.

**Expression analysis by RT-qPCR**. RNA was isolated using TRIzol reagent (Life Technologies) following the manufacturer's instructions. RNA was retrotranscribed into cDNA using PrimeScript RTPCR Kit (Clontech, Takara) and RT-qPCR was performed using SYBR Green PCR Master Mix (Applied Biosystems) in QuantStudio Real Time PCR System (Thermo Fisher Scientific), following standard protocols. See **Supplementary Table 7** for primer sequences. For statistical comparisons, normality analyses for each group were performed using Shapiro-Wilk test, followed by Levenne test for homogeneity of variances. For parametric group comparisons one-way ANOVA was used, and Kruskall-Wallis test was selected for non-parametric analyses. For multiple comparisons, Tukey correction was used for samples with homogenous variances, Tamhane's T2 for variance heterogeneity and Fisher's LSD with Bonferroni correction for comparisons among 3 groups.

**Western blot**. For all the western blot presented in this work, cell protein extracts were obtained by lysis buffer containing 1% Triton X-100, 150 mM NaCl, 50 mM Tris [pH 8], supplemented with 1x protease inhibitor cocktail (Complete Mini, Roche), 10 mM NaF and 1 mM sodium orthovanadate for 30 minutes at 4 $^{o}$C and quantified using BCA (Pierce BCA Protein Assay Kit, Thermo Fisher Scientific) or Bradford (Protein Assay Dye Reagent Concentrate, BioRad) colorimetric assays. Primary antibodies and specific conditions used were: anti-PRDM5 (sc-376277; Santa Cruz Biotechnologies; 1:2,000); anit-TXN (#2429; Cell Signalling; 1:1,000) and as a loading control anti-β-actin (A5441; Sigma; 1:5,000).

**Apoptosis assay**. Cells were harvested and stained with FITC Annexin V Apoptosis Detection Kit I (BD Biosciences) following the manufacturer's protocol at indicated time points. The percentage of Annexin $V^{+}$ cells was analyzed using FACS Canto II (BD Biosciences) and FlowJo flow cytometry analysis software. For statistical comparisons, normality analyses for each group were performed using Shapiro-Wilk test, followed by Levenne test for homogeneity of variances. For parametric group comparisons one-way ANOVA was used, and Kruskall-Wallis test was selected for non-parametric analyses. For multiple comparisons, Tukey correction was used for samples with homogenous variances, Tamhane's T2 for variance heterogeneity and Fisher's LSD with Bonferroni correction for comparisons among 3 groups.

**Cell viability assays by MTS.** Cell lines infected with viral shRNA vectors were seeded at 40,000 cells/well in 96 well plates in presence or absence of doxycycline. The number of viable cells in proliferation was determined by MTS assay (CellTiter 96 Aqueous One Solution Cell Proliferation Assay, Promega) following the manufacturer's protocol, at indicated time points. Absorbance was measured at $\lambda$ = 490nm. Cell viability in each time point was calculated as the percentage of total absorbance cells in each condition/absorbance of control cells. For statistical comparisons, normality analyses for each group were performed using Shapiro-Wilk test, followed by Levenne test for homogeneity of variances. For parametric group comparisons one-way ANOVA was used, and Kruskall-Wallis test was selected for non-parametric analyses. For multiple comparisons, Tukey correction was used for samples with homogenous variances and Tamhane's T2 for variance heterogeneity.

**Cell proliferation assays by flow cytometry.**Cell proliferation rate of MM cell lines edited by CRISPR-Cas9 system was monitored by flow cytometry. 72h after gRNA transduction, 100,000 $BFP^+$ (for single gRNAs system) or double $BFP^+$ / $GFP^+$ cells (for paired gRNAs system) were mixed with 100,000 wild type (WT, $BFP^-$ / $GFP^-$) cells. The percentage of $BFP^+$ or $BFP^+$ / $GFP^+$ cells was quantified by flow cytometry (FACS Canto II, BD Biosciences) and monitored every 72 hours during an 18-day culture. Each condition was normalized to the scramble gRNA, to assess the effect of CRISPR-Cas9 editing on cell proliferation. For statistical comparisons, normality analyses for each group were performed using Shapiro-Wilk test, followed by Levenne test for homogeneity of variances. For parametric group comparisons one-way ANOVA was used, and Kruskall-Wallis test was selected for non-parametric analyses. For multiple comparisons LSD with Bonferroni correction for comparisons among 3 groups.
