## Supplementary figures for "Chromatin activation as a unifying principle underlying pathogenic mechanisms in multiple myeloma"

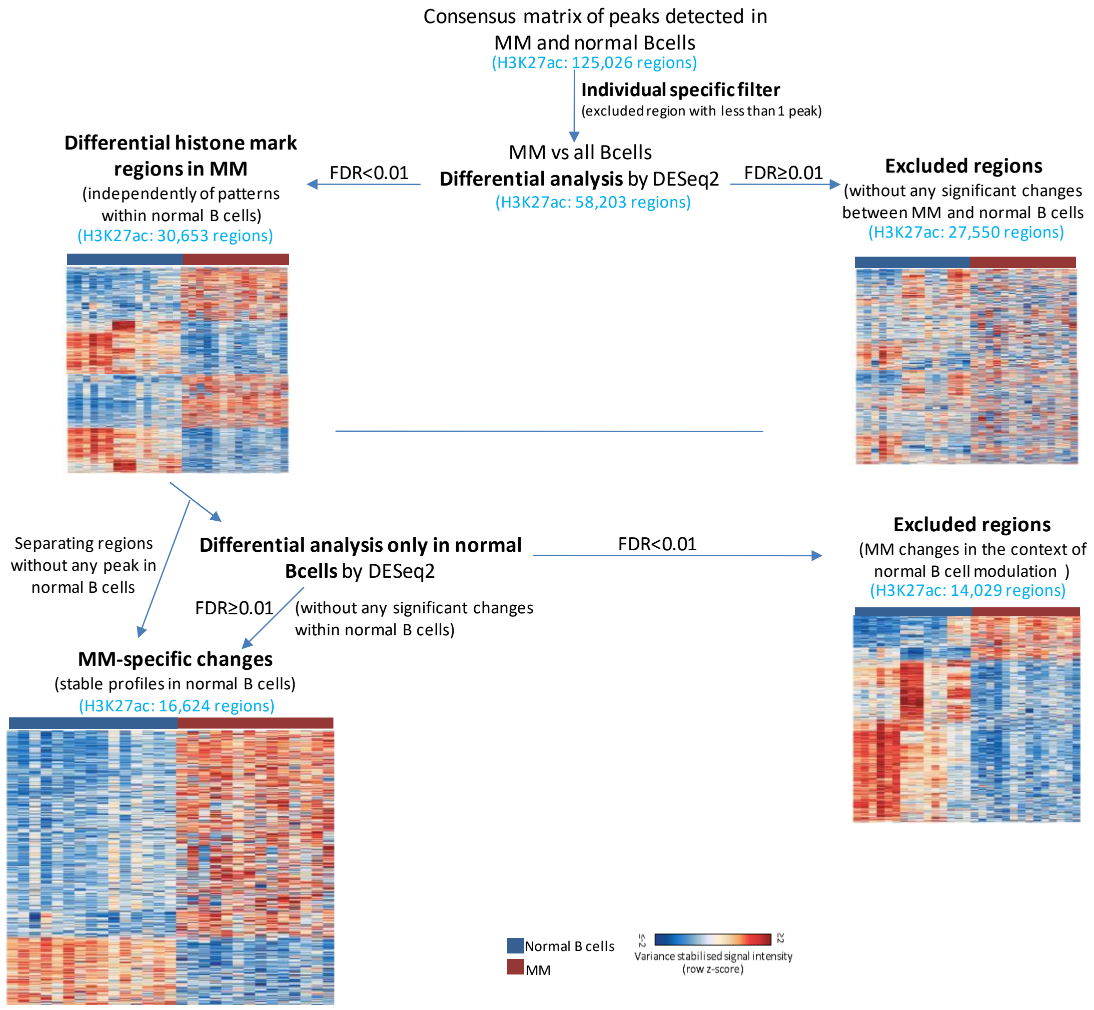


**Supplementary Figure 1.** Strategy of differential histone marks analysis in MM as compared to normal B cells. As a final step we segregated the regions into those with either dynamic (lower left heatmap) or stable (lower right heatmap) patterns in normal B cell differentiation.

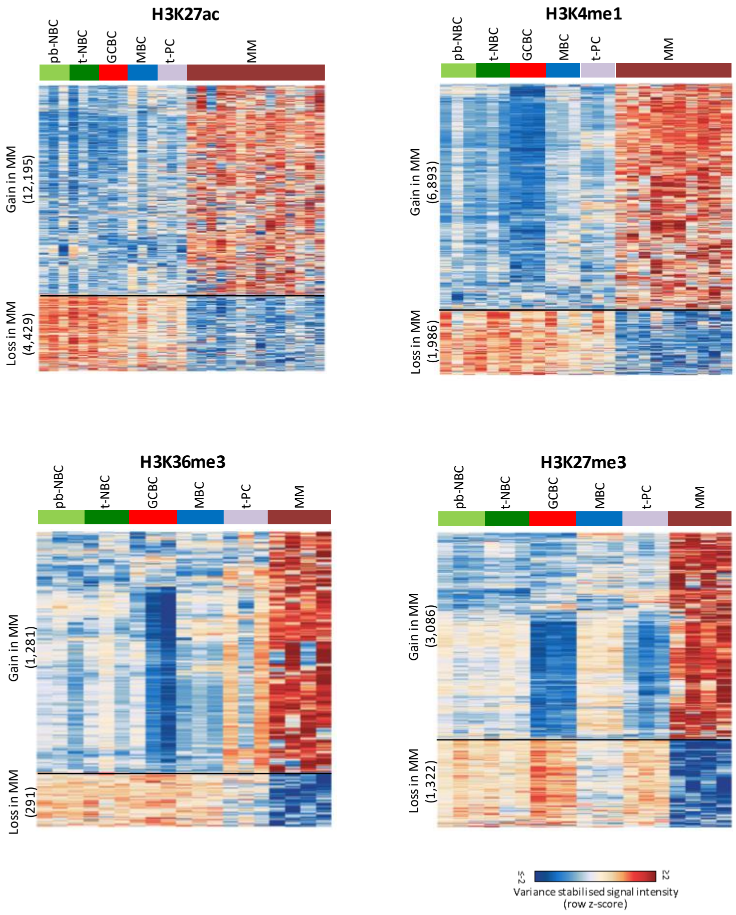


**Supplementary Figure 2.** Heatmaps representing histone mark changes specifically acquired in MM as compared to normal B cells with stable chromatin profiles throughout cell differentiation. The upper part of each heatmap represent the regions with increase of a particular histone mark in MM (i.e. gain in MM), while the lower part represents the regions with decrease in MM (i.e. loss in MM). Number of regions are in brackets. pb-NBC, naive B cells from blood; t-NBC, naive B cells from tonsils; GCBC, germinal centre B cells; MBC, memory B cells; t-PC, plasma cell from tonsils; MM, multiple myeloma.

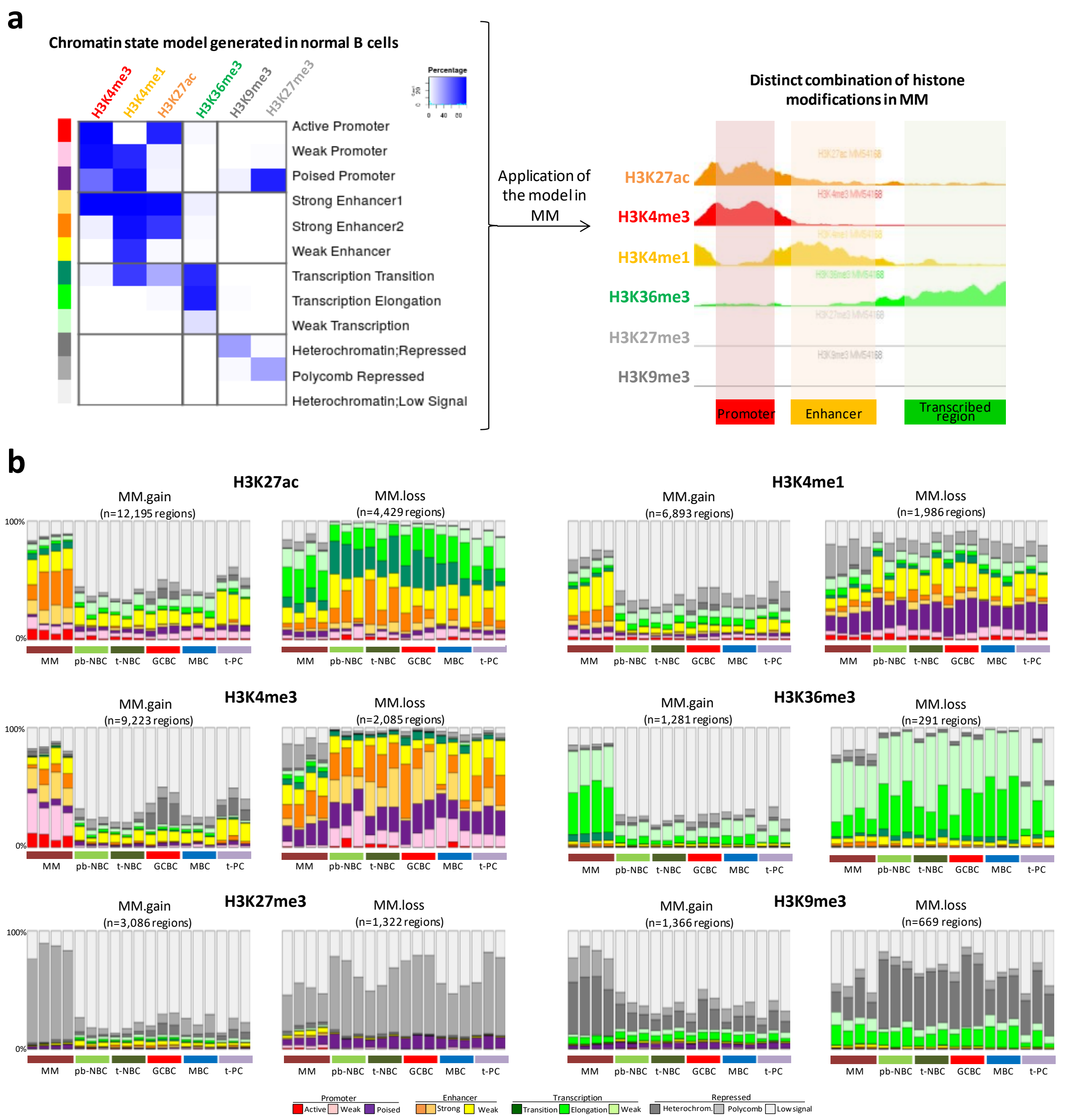


**Supplementary Figure 3.** **a)** Schematic representation of chromatin states generation strategy. Left panel shows emissions of the chromatin state model generated in normal B cells[11], where the percentages of regions assigned to a specific chromatin state (rows) that contain a specific histone mark (columns). Right panel shows how this model was applied to our MM data. **b)** Distribution of the different chromatin states in all analyzed samples separately at regions with differential enrichment of histone marks in MM (either gain or loss) as compared to stable chromatin profiles throughout B cell differentiation. pb-NBC, naive B cells from blood; t-NBC, naive B cells from tonsils; GCBC, germinal centre B cells; MBC, memory B cells; t-PC, plasma cell from tonsils; MM, multiple myeloma.

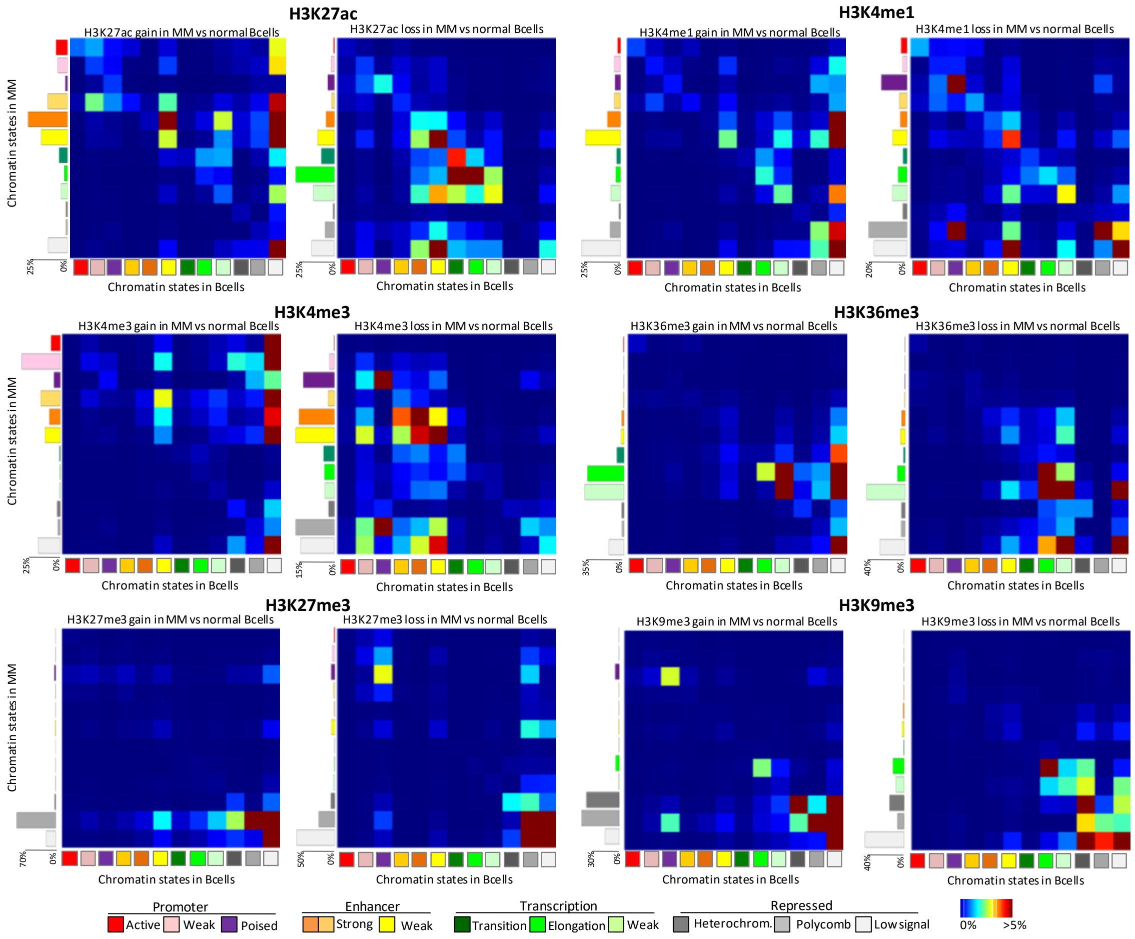


**Supplementary Figure 4.** Chromatin state transition matrix for regions with differential enrichment of six histone marks in MM (either gain or loss) as compared to normal B cells. Columns represent the chromatin state in normal B cells and rows are chromatin states in MMs that arise from normal B cells. The total matrix represents 100 percent of the differential regions. The percentage of regions associated with each chromatin state in MM patients is indicated in the left side bar chart.

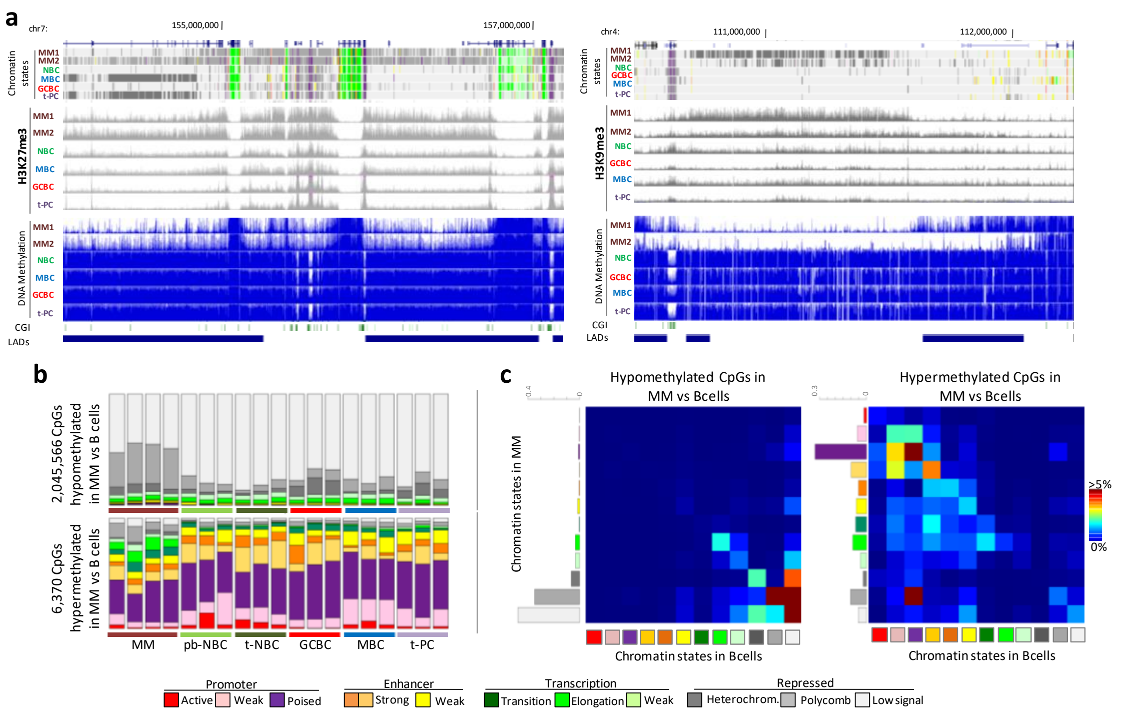

**Supplementary Figure 5. a)** Schematic representation of large blocks, with increased peaks of H3K27me3 (left) and H3K9me3 (right) in MM, harbouring extensive loss of DNA methylation in neoplastic cells, as compared to normal B cell differentiation. For each panel following tracks, showing different epigenetic features in the very same samples, are represented (from upper to lower): chromatin states segmentation; H3K27me3 (left) or H3K9me3 (right) peaks profile and DNA methylation values (0-1 scale). Additionally tracks showing CpG Island (CGI) and lamin-associated domains (LADs, data from fibroblasts by ENCODE) are added. We detected 26,537 of these large blocks of DNA hypomethylated regions (DNA methylation valleys, range size 5kb-40kb). **b)** Distribution of the different chromatin states in all analyzed samples separately at CpG sites hypomethylated or hypermethylated in MM as compared to normal B cell differentiation. **c)** Chromatin state transition matrix for CpG sites hypomethylated or hypermethylated in MM as compared to normal B cell differentiation. Columns represent the chromatin state in normal B cells and rows are chromatin states in MMs that arise from normal B cells. The total matrix represents 100 percent of the differential regions. The percentage of regions associated with each chromatin state in MM patients is indicated in the left side bar chart. pb-NBC, naive B cells from blood; t-NBC, naive B cells from tonsils; GCBC, germinal centre B cells; MBC, memory B cells; t-PC, plasma cell from tonsils; MM, multiple myeloma.

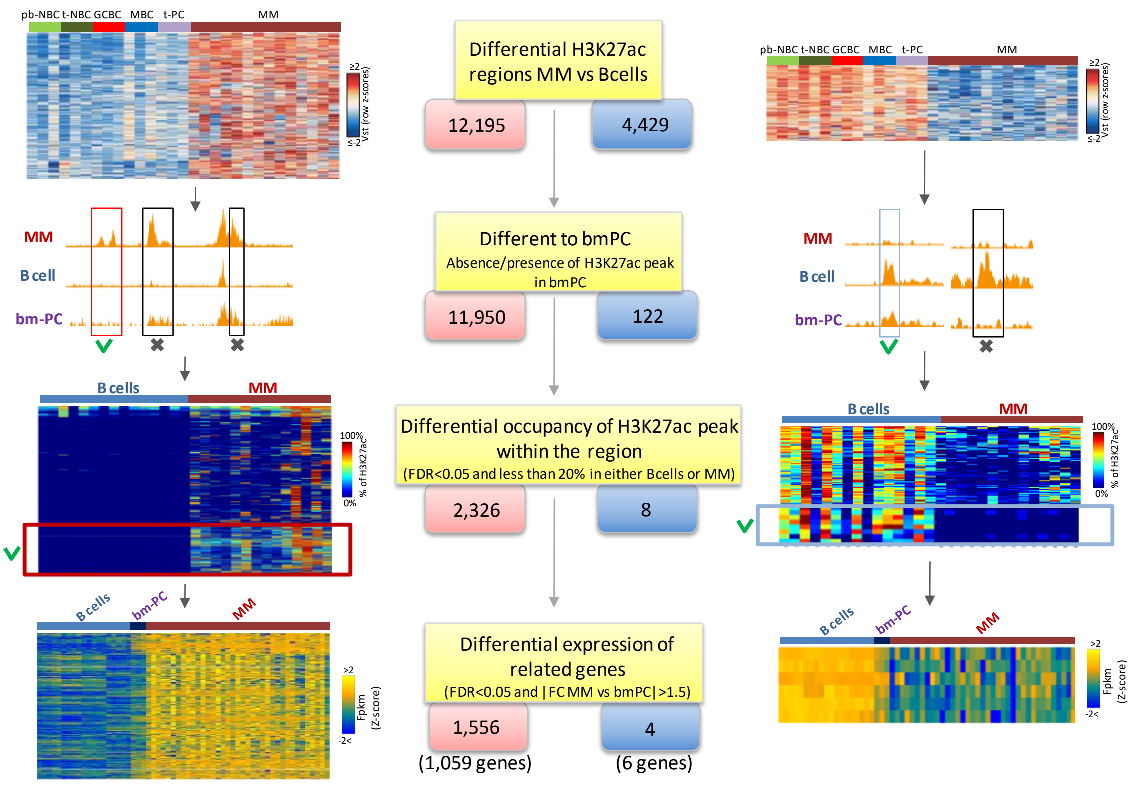


**Supplementary Figure 6.** Schematic representation of strategy to determine *de novo* active regions in MM, that is those without any H3K27ac peak across normal B cell differentiation including bm-PCs (i.e. inactive in normal B cells), while gaining this histone mark only in MM (numbers in red boxes and left panels). An opposite analysis in order to detect *de novo* inactive regions in MM was performed in parallel (numbers in blue boxes and right panels). For a detailed explanation of each step please refer to Methods section. Graphical representation of regions selected in each step is shown on left (for regions *de novo* active in MM) and right (for regions *de novo* inactive in MM) from the schematic workflow. pb-NBC, naive B cells from blood; t-NBC, naive B cells from tonsils; GCBC, germinal centre B cells; MBC, memory B cells; t-PC, plasma cell from tonsils; bm-PC, plasma cell from bone marrow; MM, multiple myeloma.

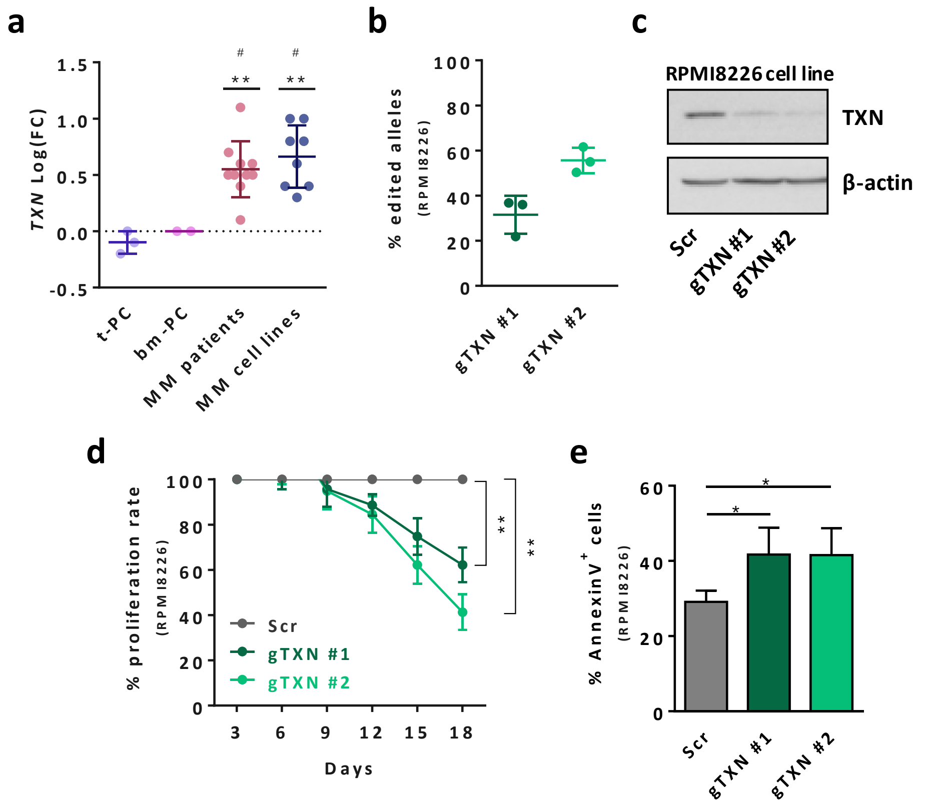


**Supplementary Figure 7.a**) *TXN* overexpression validation in an independent cohort of MM patients and cell lines versus t-PC (* p <0.05, ** p < 0.01, *** p < 0.001, ns: not significant) and bm-PC ($^{\#}$ p <0.05, $^{\#\#}$ p < 0.01, $^{\#\#\#}$ p < 0.001, ns: not significant) as determined by RT-qPCR. **b)** Estimation of the allelic cell population percentage exhibiting indel events in the targeted site of the RPMI8226 cell line, analyzed by Tracking of Indels by DEcomposition (TIDE) web tool. **c)** Validation of TXN knockout by western blot analysis in the RPMI8226 cell line. **d)** Cell proliferation rates of scramble cells (Scr) and cells harboring two different gRNAs in RPMI8226 cell line, as determined by flow cytometry analysis. **e)** Effect of TXN knockout on cell apoptosis in the RPMI8226 cell line, as determined by Annexin V flow cytometry analysis. * p<0.05, ** p<0.01, *** p<0.001, ns: not significant.

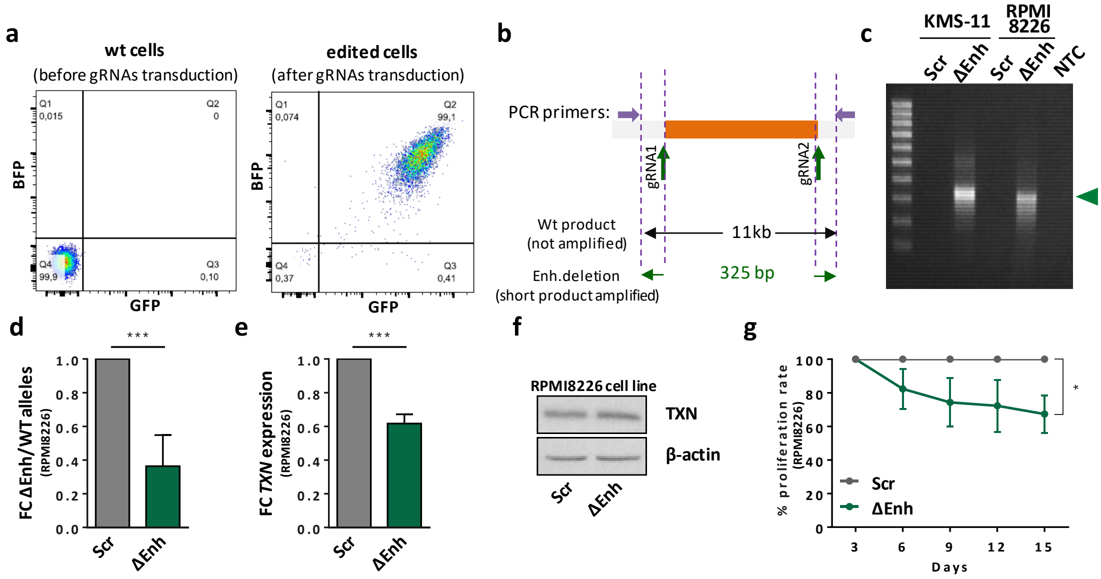


**Supplementary Figure 8. a)** Fraction of double positive BFP⁺/GFP⁺ cells analyzed by flow cytometry prior to (left) and after paired gRNAs transduction (right). Upper right quadrant (Q2) shows double BFP⁺ / GFP⁺ cells. The cell number proportion of each subpopulation is shown in each quadrant. **b)** Primer design strategy for qualitative analysis of *TXN* enhancer deletion. External primers flanking the enhancer target region (11 kb) were designed to detect the presence of smaller PCR products (325 bp) in edited alleles by gel electrophoresis. **c)** Qualitative analysis of *TXN* enhancer deletion by electrophoresis of genomic DNA PCR products. Wild type genomic DNA (scramble cells, Scr) and water (NTC) templates are used as negative controls. The green arrowhead indicates the size of PCR products expected from the deleted allele. **d)** Quantification of *TXN* enhancer deletion in the RPMI8226 cell line by genomic DNA qPCR normalized to a distal non-targeted genomic region, represented as fold change of deleted enhancer (ΔEnh) versus wild type (WT) alleles. **e)** *TXN* mRNA expression levels determined by RT-qPCR in the RPMI8226 cell line. **f)** TXN protein levels determined by western blot in the RPMI8226 cell line. **g)** Cell proliferation rates of Scr cells and cells harboring the enhancer deletion in RPMI8226 cell line, as determined by flow cytometry analysis. * $p<0.05$, ** $p<0.01$, *** $p<0.001$, ns: not significant.

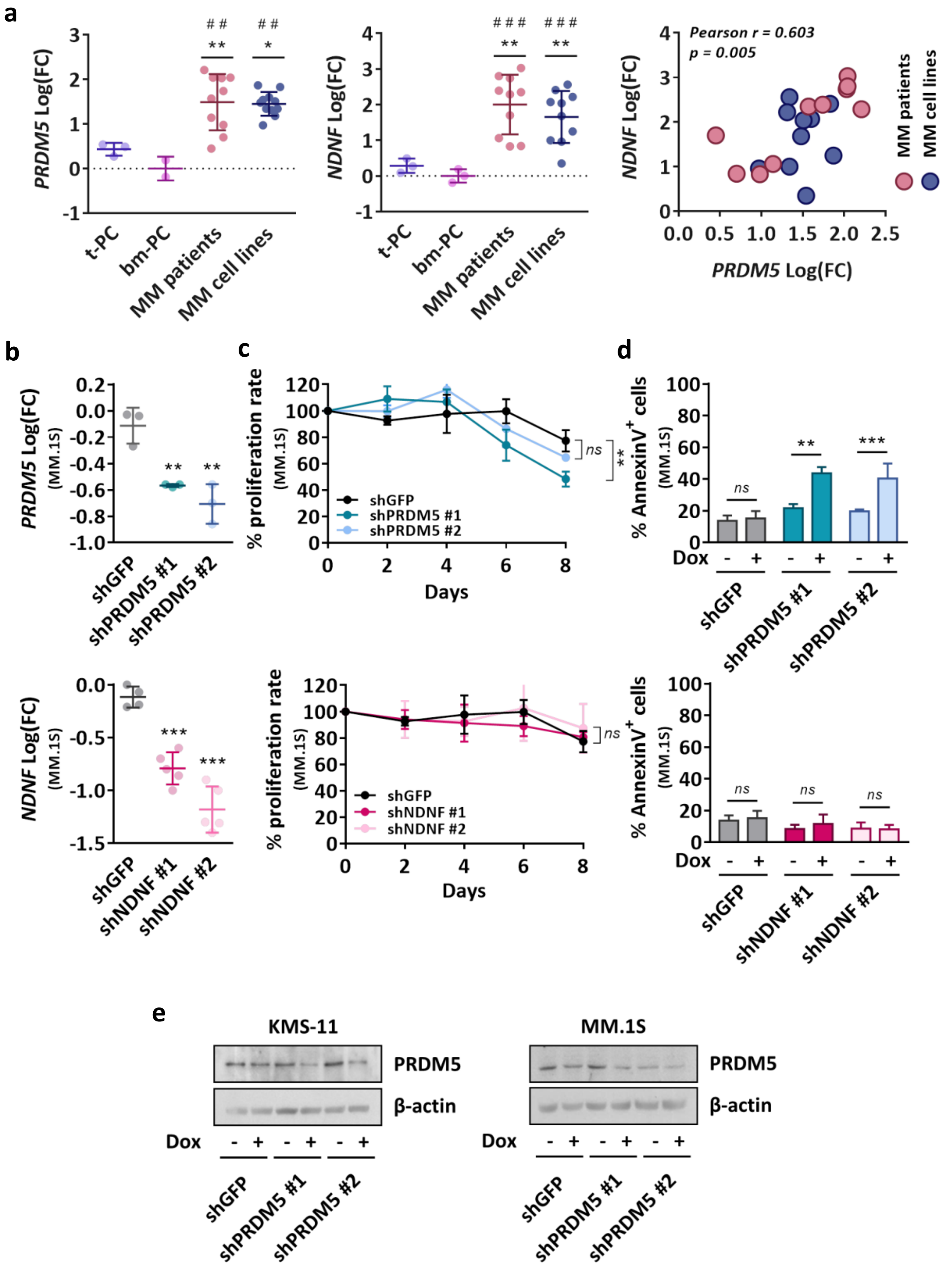


**Supplementary Figure 9. a)** *PRDM5* and *NDNF* overexpression validation in an independent cohort of MM patients and cell lines versus t-PC (* p<0.05, ** p<0.01, *** p<0.001, ns: not significant) and bm-PC (# p<0.05, ## p<0.01, ### p<0.001, ns: not significant) as determined by RT-qPCR, and correlation of expression levels of both transcripts (right panel). **b)** Validation of *PRDM5* (top) and *NDNF* (bottom) gene knockdown after shRNA expression determined by RT-qPCR in the MM.1S cell line. Expression values are normalized to the same condition prior to addition of doxycycline. Statistical analysis compares the effect of each shRNA versus the scramble shRNA (shGFP). **c)** Relative cell proliferation rate (%) of the MM.1S cell line normalized to the same condition prior to addition of doxycycline. Upper panel present the results for *PRDM5* knockdown cells and lower panel for *NDNF* knockdown cells. Statistical analysis compares the effect of each shRNA versus the scramble shRNA (shGFP). **d)** Effect of *PRDM5* (upper panel) and *NDNF* (lower panel) knockdown in cell apoptosis, as determined by Annexin V flow cytometry analysis in the MM.1S cell line. **e)** Reduction of PRDM5 protein levels after shRNA induction determined by western blot in the KMS-11 and MM.1S cell lines. Dox: Doxycycline. * $p<0.05$, ** $p<0.01$, *** $p<0.001$, ns: not significant.

### Supplementary Tables

**Separately provided tables:**

**Supplementary Table 1. Experimental design**. MM and normal B cell cases used for the study

**Supplementary Table 2. Characterization of Multiple Myeloma (MM) patient samples**.

**Supplementary Table 3. List of de novo activated regions in MM with associated target genes.**

**Supplementary Table 4. TFBS enrichment analysis by MEME.** TF motifs analysis in 806 sites with increased chromatin accessibility in MM, within the 1,556 regions de novo active in MM.

**Supplementary Table 5. Expression of all members of IRF, FOX and MEF2 transcription factor families.** FPKM values from RNA-Seq of additional MM and normal B cell series are shown.

**Supplementary Table 6. Genes included in co-regulated chromatin regions**. Gene pair selected for downstream analysis is shaded.

**Supplementary Table 7. Primers and target sequences used in this study**
